# UTX coordinates TCF1 and STAT3 to control progenitor CD8^+^ T cell fate in autoimmune diabetes

**DOI:** 10.1101/2024.12.12.628206

**Authors:** Ho-Chung Chen, Madison F. Bang, Hsing-Hui Wang, Karl B. Shpargel, Lisa A. Kohn, David Sailer, Shile Zhang, Ethan C. McCarthy, Maryamsadat Seyedsadr, Satchel Stevens, Caitlyn L. H. Pham, Zikang Zhou, Xihui Yin, Nicole M. Wilkinson, Esther M. Peluso, Christian Bustillos, Jessica G. Ortega, Lixin Yang, Ashlyn A. Buzzelli, Reina Capati, Dennis J. Chia, Steven D. Mittelman, Christina M. Reh, Jason K. Whitmire, Melissa G. Lechner, Willy Hugo, Maureen A. Su

## Abstract

Type 1 Diabetes Mellitus (T1D) is a chronic disease caused by an unremitting autoimmune attack on pancreatic beta cells. This autoimmune chronicity is mediated by stem-like progenitor CD8^+^ T cells that continually repopulate the pool of beta cell-specific cytolytic effectors. Factors governing the conversion of progenitors to effectors, however, remain unclear. T1D has been linked to a chromosomal region (Xp13-p11) that contains the epigenetic regulator UTX, which suggests a key role for UTX in T1D pathogenesis. Here, we show that T cell-specific UTX deletion in NOD mice protects against T1D development. In T cells of NOD mice and T1D patients, UTX ablation resulted in the accumulation of CD8^+^ progenitor cells with concomitant deficiency of effectors, suggesting a key role for UTX in poising progenitors for transition to effectors. Mechanistically, UTX’s role in T1D was independent of its inherent histone demethylase activity but instead relied on binding with transcription factors (TCF1 and STAT3) to co-regulate genes important in the maintenance and differentiation of progenitor CD8^+^ T cells. Together, these findings identify a critical role for UTX in T1D and the UTX:TCF1:STAT3 complex as a therapeutic target for terminating the long-lived autoimmune response.

## INTRODUCTION

Type 1 Diabetes (T1D) is a chronic autoimmune disease in which insulin-producing beta cells are destroyed by a sustained T cell attack (1). As a consequence, clinical symptoms do not remit, and patients require lifelong exogenous insulin administration. This contrasts with Guillain Barre Syndrome, acute disseminated encephalomyelitis, and other self-limited autoimmune conditions in which the immune response wanes and clinical symptoms usually resolve within months. What underlies autoimmune persistence in T1D is not completely understood and identifying mechanisms that perpetuate chronic immune-mediated destruction is an important step toward developing therapeutic approaches that interrupt this process.

Much has been learned about the mechanisms of immune chronicity through studies in models of chronic virus infection and progressive cancer. Similar to the T1D autoimmune response, the immune responses in chronic virus infection and progressive cancer are characterized by continued T cell activation by persistent antigens and a prominent, long-lived CD8^+^ T cell stem-like progenitor (T_prog_) population (2–4). In all three scenarios, these T_prog_ cells sustain persistent immune responses by continually replenishing the pool of terminally differentiated CD8^+^ T cells. In chronic virus infection and progressive cancers, T_prog_ cells give rise to CD8^+^ T cell effector populations that become exhausted and fail to eradicate the immune threat (5, 6). In T1D, on the other hand, these short-lived, terminally differentiated CD8^+^ T mediator cells (T_med_) retain their cytolytic function and aggressively destroy beta cells (3). Thus, a critical feature of T1D pathogenesis is the persistent conversion of long-lived CD8^+^ T_prog_ cells to T_med_ cells with effector function.

Through genetic linkage studies, T1D has been associated with gene loci with key roles in T_prog_ cells. For instance, T1D has been associated with polymorphisms in the gene encoding TCF1 (7, 8), a transcription factor critical in T_prog_ cell formation, maintenance and function (9). Additionally, accelerated T1D in mice and humans has been linked with gain-of-function mutations in STAT3 (10), a transcription factor implicated in the conversion of T_prog_ to effector (11). Remarkably, T1D acceleration in STAT3 gain-of-function mice has been attributed to increased CD8+ T cell effectors (10, 12), a finding in line with STAT3’s role in CD8^+^ T_prog_ to effector conversion (11). Together, these findings highlight the importance of genetic regulators of T_prog_ cells in the T1D autoimmune response.

T1D has also been linked to a region of chromosome X (Xp13-p11) that contains the epigenetic regulator UTX (Ubiquitously transcribed tetratricopeptide repeat, X chromosome; encoded by *KDM6A*) (13). UTX has pleiotropic functions, including intrinsic histone demethylase activity (14, 15). Whether UTX plays a role in T1D, and the cellular and molecular mechanisms by which UTX may function, however, remain unclear (13). Here, we show that T cell-specific UTX expression is required for T1D through its role in driving stem-like T_prog_ cells to become cytolytic T_med_ cells. This role is independent of its demethylase function but instead involves cooperative interactions between UTX and the transcription factors TCF1 and STAT3. UTX, TCF1, and STAT3 co-occupancy at key progenitor and effector gene loci enforces a transcriptional program that drives T_prog_ to effector conversion. These findings point to UTX as a critical regulator of the T1D autoimmune response and point to targeting UTX: TCF1: STAT3 mechanisms as a potential approach for interrupting the persistent autoimmune attack.

## RESULTS

### UTX in T cells of NOD mice is required for T1D and alters CD8^+^ T cell subset distribution

UTX has been implicated in controlling T cell differentiation in anti-cancer and anti-viral immunity (16–18), but its role in diabetogenic T cell responses is unknown. Given genetic data linking a UTX-containing chromosomal region and T1D, we hypothesized that UTX may play a key role in T cell-mediated T1D pathogenesis. To test this, we generated NOD mice with T cell-specific UTX deletion (*UTX^fl/fl^ Lck-Cre^+^,* herein referred to as *NOD-UTX^TCD^*). As expected, UTX protein was decreased in T cells isolated from *NOD-UTX^TCD^* mice, compared to UTX-sufficient littermate controls (*UTX^fl/fl^ Lck-Cre*^-^; herein referred to as *NOD-WT*) **(Supplemental Figure 1A)**. By 30 weeks of age, none of the *NOD-UTX^TCD^* female mice developed diabetes, in contrast to almost 80% of *NOD-WT* female littermate controls (**Figure 1A**). Hematoxylin/eosin (H/E) stained pancreas sections showed significantly reduced immune infiltration in the islets of *NOD-UTX^TCD^* mice, compared to *NOD-WT* controls (**Figure 1, B and C**). Notably, this protection from diabetes is independent of UTX’s intrinsic histone demethylase activity, since NOD mice with point mutations in UTX’s demethylase domain (*NOD-UTX^Demethylase-dead^* or *NOD-UTX^DMD^* mice) continue to develop diabetes at the same rate as *NOD.WT* controls **(Supplemental Figure 1B)**. Thus, T cell-specific UTX deficiency protects against diabetes and insulitis in NOD mice in a demethylase-independent manner.

**Figure 1.**
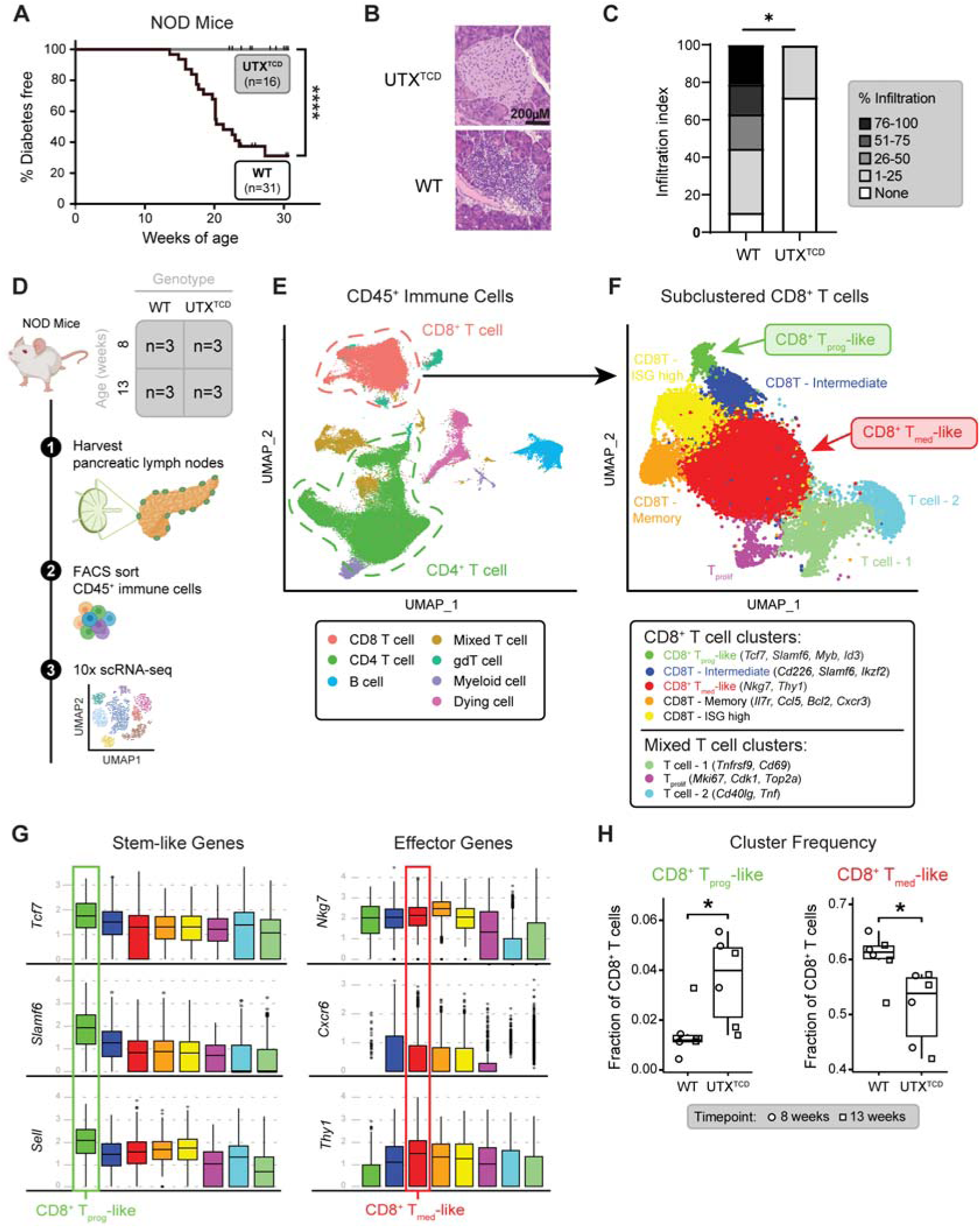
Resistance to T1D in *NOD-UTX^TCD^* mice is associated with altered distribution of pLN CD8^+^ T cells. **(A)** Diabetes-free incidence curves of *NOD-UTX^TCD^* vs. *NOD-WT* female littermates. ****p<0.0001; Log rank test. **(B and C)** Representative H/E stained pancreas sections (B) and cumulative islet infiltration scores (C) from *NOD-UTX^TCD^* vs. *NOD-WT* female littermates. *p<0.05; Tukey’s multiple comparisons test. **(D)** Scheme for 10X genomics single-cell RNA sequencing (scRNAseq) of CD45^+^ cells from pLN of *NOD-WT* and *NOD-UTX^TCD^* mice. **(E and F)** UMAP of CD45^+^ cells (E) and subclustered CD8^+^ T cells (F). **(G)** Gene expression across the CD8^+^ T cell clusters. **(H)** Comparison of subset frequencies of CD8^+^ T_prog_-like and CD8^+^ T_med_-like populations. CD8^+^ T_prog_-like, *p=0.015; CD8^+^ T_med_-like, *p=0.026; two-sided, unpaired Mann-Whitney test.

We next sought to explore the cellular mechanism by which UTX functions in T cells to promote T1D. Prior to infiltrating pancreatic islets, diabetogenic T cells are first primed in the pancreatic lymph nodes (pLN) (3, 19). To delineate how UTX deficiency alters pLN T cell dynamics in NOD mice, we performed single-cell RNA seq (scRNAseq) of pLN immune cells (CD45^+^) from 8 and 13-week-old *NOD-WT* and *NOD*-*UTX^TCD^* mice (**Figure 1D**; n=6 per genotype). These ages represent distinct stages in T1D autoimmunity development: 8 weeks represents a stage when immune cell infiltration is prominent in islets, and 13 weeks represents a stage immediately prior to overt diabetes (20). UMAP clustering of 97,400 cells showed 7 major immune cell populations including CD8^+^ T cells, CD4^+^ T cells, B cells, and myeloid cells (**Figure 1E and Data File S1).** Frequencies of these populations were similar between *NOD-WT* vs. *NOD-UTX^TCD^* mice **(Supplemental Figure 1, C and D).** We further evaluated CD4^+^ and CD8^+^ T cell populations to assess for primary alterations resulting from T cell-specific UTX deficiency.

Sub-clustering of CD4^+^ T cells yielded 10 populations **(Supplemental Figure 2A)**, including T follicular helper (Tfh) and regulatory T (Treg) cells. Frequencies of key autoimmunity-associated CD4^+^ T cell subsets in the pLN were similar between *NOD-WT* vs. *NOD-UTX^TCD^* mice **(Supplemental Figure 2B)**. Moreover, flow cytometric analysis of pLN showed similar frequencies of CD4^+^ T cell subsets [T follicular helper (Tfh; CXCR5^+^ PD1^+^), T helper 1 (Th1; T-bet^+^), and regulatory T cells (Treg; Foxp3^+^)] **(Supplemental Figure 2C)** and pro-inflammatory cytokine-producing CD4^+^ cells **(Supplemental Figure 2D)** between *NOD-WT* vs. *NOD-UTX^TCD^* mice. Thus, pLN CD4^+^ T cell distribution was largely unchanged by T cell-specific UTX deficiency in NOD mice.

In contrast, significant changes were noted in the distribution of CD8^+^ T cell subsets between *NOD-WT* and *NOD-UTX^TCD^*mice. CD8^+^ T cells were sub-clustered into 8 populations (**Figure 1F**), which included a “CD8^+^ T_prog_-like” population that expressed genes associated with stem-like progenitor cells (*Tcf7* [which encodes TCF1]*, Slamf6* [which encodes Ly108]*, Sell, Myb, Ikzf2, Id3*) (**Figure 1G and Supplemental Figure 3A)** and a “CD8^+^ T_med_-like” effector population that expressed genes associated with terminally-differentiated autoimmune mediators (*Nkg7, Cxcr6, Thy1*) (3, 9) (**Figure 1G**). Strikingly, a higher frequency of CD8^+^ T_prog_-like, and a lower frequency of CD8^+^ T_med_-like, cells were seen in the pLN of *NOD-UTX^TCD^* mice, compared to *NOD-WT* littermates (**Figure 1H and Supplemental Figure 3, B and C).** Thus, protection from T1D in *NOD-UTX^TCD^*mice is associated with an altered distribution of pLN CD8^+^ T cell subsets expressing markers of stem-like progenitors and effectors.

### UTX in T cells promotes CD8^+^ T cell transition from progenitor to cytolytic effector state in T1D

Within the pLN of NOD mice, stem-like progenitors (T_prog_) have been reported to differentiate into terminally-differentiated autoimmune mediators (T_med_), which then enter the pancreatic islets to destroy beta cells (3). In line with this report, pseudotime trajectory analysis of our scRNAseq dataset suggested a differentiation pathway that starts with the CD8^+^ T_prog_-like population, moves through an intermediate population (CD8^+^ T Intermediate), and ends in the CD8^+^ T_med_-like population (**Figure 2A**). Along the pseudotime axis, expression of stem-like progenitor-associated genes (*Tcf7*, *Slamf6*) was highest in CD8^+^ T_prog_-like, decreased in the CD8^+^ T Intermediate, and lowest in CD8^+^ T_med_-like cells (**Figure 2, B and C).** On the other hand, expression of effector-associated genes (*Thy1, Cxcr6*) was lowest in the CD8^+^ T_prog_-like, and increased in CD8^+^ T-Intermediates and CD8^+^ T_med_-like cells (**Figure 2, B and C)**. Of note, T cell-specific UTX deficiency altered this differentiation trajectory. Comparison of cell densities along the pseudotime trajectory revealed relative accumulation of *NOD-UTX^TCD^*CD8^+^ T_prog_-like cells, compared to *NOD-WT* (**Figure 2D**). This difference was accompanied by a relative decrease in *NOD-UTX^TCD^*CD8^+^ T_med_-like cells, compared to *NOD-WT*. Together, these data suggest a potential block in the conversion of CD8^+^ T_prog_ to T_med_ cells in *NOD-UTX^TCD^*mice.

**Figure 2.**
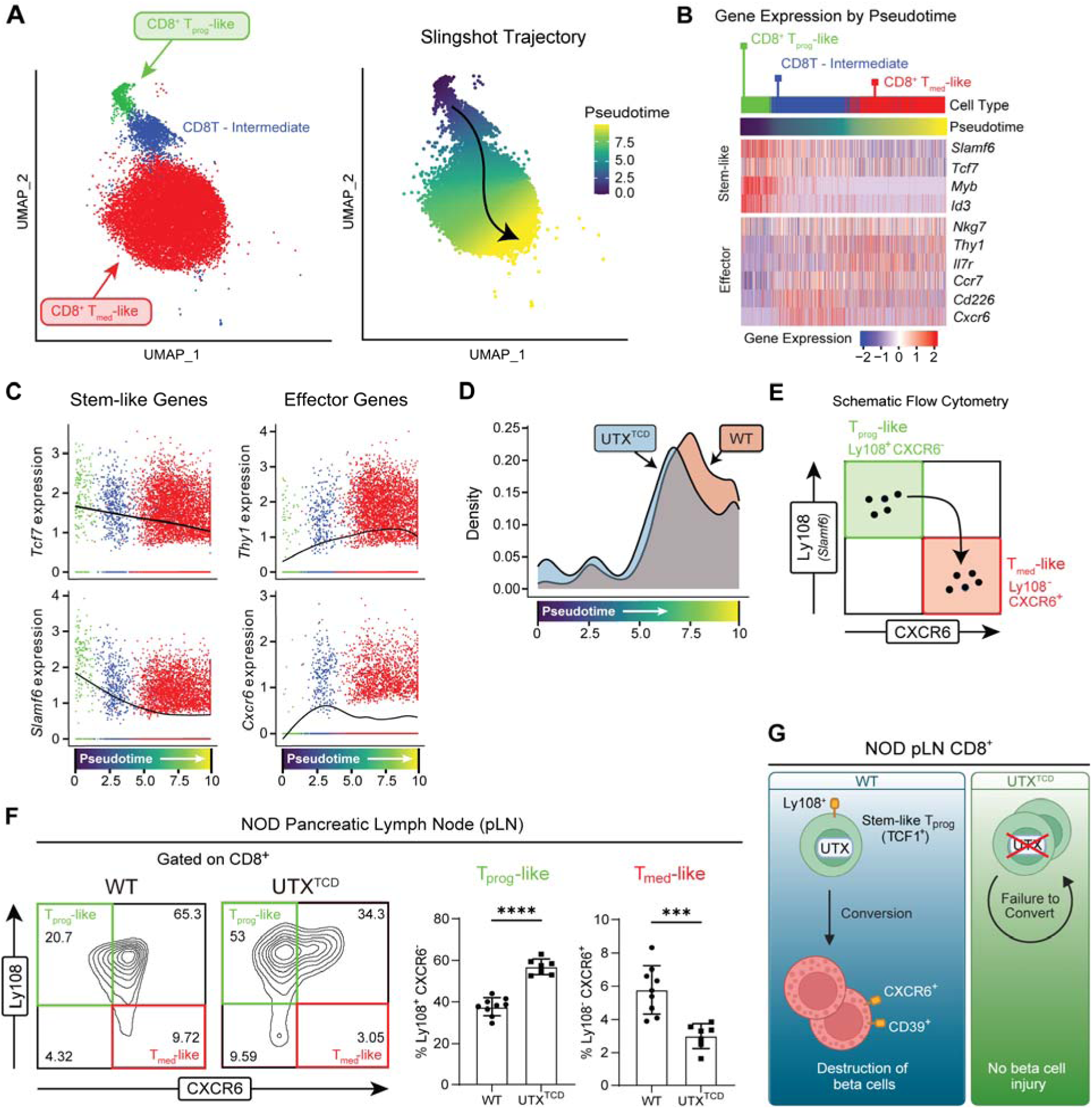
Impaired conversion of stem-like progenitors to effectors in *NOD-UTX^TCD^* mice. **(A)** UMAP plot of CD8^+^ T_prog_-like, CD8^+^ T Intermediate, and CD8^+^ T_med_-like clusters (left), with cells color-coded chronologically along Slingshot pseudotime (right). **(B)** Heatmap of scRNAseq expression for key genes, plotted along Slingshot pseudotime. **(C)** Expression of key genes along Slingshot pseudotime trajectory, color coded by clusters shown in UMAP (A, left). **(D**) Comparison of cell densities of CD8^+^ T_prog_-like, CD8^+^ T Intermediate, and CD8^+^ T_med_-like cells derived from 8-week-old *NOD-WT* (WT) vs. *NOD-UTX^TCD^* (UTX^TCD^) mice, plotted along Slingshot pseudotime trajectory. P < 2.2 e-16; Asymptotic two-sample Kolmogorov-Smirnov test. (**E)** Gating scheme for identifying T_prog_-like and T_med_-like cells by flow cytometric analysis. **(F)** Representative flow cytometric plot (left) and average frequencies (right) of pLN T_prog_-like (Ly108^+^ CXCR6^-^) and T_med_-like (Ly108^-^ CXCR6^+^) cells of *NOD*-*WT* and *NOD-UTX*^TCD^ female littermates (15-29 weeks of age). ****p<0.0001;***p<0.001; Student’s t test. (**G)** Working model: UTX promotes conversion of long-lived stem-like CD8^+^ T cell progenitors (T_prog_) to short-lived mediators (T_med_), which then enter pancreatic islets to mediate beta cell destruction.

To verify this finding, we turned to flow cytometric analysis, using previously reported cell surface markers to identify T_prog_-like and T_med_-like cells (4). In this approach, reciprocal expression of Ly108 (*Slamf6*) and CXCR6 were used to distinguish T_prog_-like (Ly108^+^ CXCR6^-^) and T_med_-like (Ly108^-^ CXCR6^+^) populations (**Figure 2E**). Supporting the validity of this gating strategy, expression of the stem-like transcription factor TCF1, was higher in Ly108^+^ CXCR6^-^T_prog_-like than Ly108^-^ CXCR6^+^ T_med_-like populations **(Supplemental Figure 4A).** Further, expression of the CD39 mediator marker was lower in Ly108^+^ CXCR6^-^ T_prog_-like cells than Ly108^-^ CXCR6^+^ T_med_-like populations (**Supplemental Figure 4B).** Using this gating strategy, a significant increase in T_prog_-like cells was seen in pLN of *NOD-UTX^TCD^* mice, compared to *NOD-WT* (**Figure 2F**). At the same time, a significant decrease in T_med_-like populations was seen in pLN of *NOD-UTX^TCD^* mice, compared to *NOD-WT.* Taken together, our scRNAseq and flow cytometric analyses support a model in which UTX promotes conversion of CD8^+^ T cell progenitors to effectors, which then enter the pancreatic islets to destroy beta cells and incite diabetes (**Figure 2G**). In support of the importance of UTX in diabetogenic CD8^+^ T cells, diabetes development was delayed in immunodeficient hosts after adoptive transfer with UTX-deficient CD8^+^ T cells, compared to UTX-sufficient (wildtype) CD8^+^ T cells **(Supplemental Figure 4C)**.

### UTX controls progenitor to cytolytic effector differentiation in beta cell-specific CD8^+^ T cells

CD8^+^ T cells recognizing the pancreatic beta cell antigen IGRP (islet-specific glucose-6-phosphatase catalytic subunit-related protein) are a major pathogenic population in mice and humans with T1D (21, 22). Remarkably, Gearty et al. reported that IGRP-specific stem-like progenitors (T_prog_; Ly108^+^ CD39^-^) give rise to both T_prog_ and autoimmune mediator (T_med_; Ly108^-^CD39^+^) populations, and as few as 20 pLN T_prog_ cells from NOD donors can induce diabetes in immunodeficient hosts (3). Given their potent diabetes-inducing potential, we sought to define the role of UTX in controlling the differentiation of IGRP-specific T_prog_ cells into cytolytic T_med_.

In keeping with the markers used by Gearty et al., we identified pLN CD8^+^ T_prog_ cells as Ly108^+^ CD39^-^ cells and T_med_ cells as Ly108^-^ CD39^+^ cells (3) (**Figure 3A**). As expected, Ly108^+^ CD39^-^T_prog_ cells expressed higher levels of the progenitor marker TCF1 **(Supplemental Figure 4D)** and lower levels of the mediator marker CXCR6 than Ly108^-^ CD39^+^ T_med_ cells (**Figure 3B**). We further identified IGRP-specific T cells by their expression of CD44, a marker of antigen-experienced T cells, and their binding to tetramers containing the IGRP mimotope NRP-V7 (CD44^+^ NRP-V7^+^) (23). Using this gating approach, the frequency of T_prog_ cells among IGRP-specific, antigen-experienced CD8+ T cells (Ly108^+^ CD39^-^ cells within CD8^+^ CD44^+^ NRP-V7^+^) was increased in *NOD-UTX^TCD^* pLN compared to *NOD-WT* (**Figure 3C**). At the same time, the frequency of T_med_ (Ly108^-^ CD39^+^) within antigen-experienced, IGRP-specific (CD8^+^ CD44^+^ NRP-V7^+^) cells was significantly lower in *NOD-UTX^TCD^* pLN, compared to *NOD-WT* (**Figure 3C**). Similar findings were seen when CXCR6, rather than CD39, was used as a mediator marker **(Supplemental Figure 4E)**. With age, the frequency of IGRP-specific T_med_ (Ly108^-^ CD39^+^ cells within CD8^+^ CD44^+^ NRP-V7^+^) increased in *NOD-WT* pLN, while IGRP-specific T_prog_ (Ly108^+^ CD39^-^ cells within CD8^+^ CD44^+^ NRP-V7^+^) accumulated in *NOD-UTX^TCD^*pLN **(Supplemental Figure 4F)**. Together, these findings demonstrate a role for UTX in promoting the conversion of beta cell-specific T_prog_ to T_med_ in the pLN.

**Figure 3.**
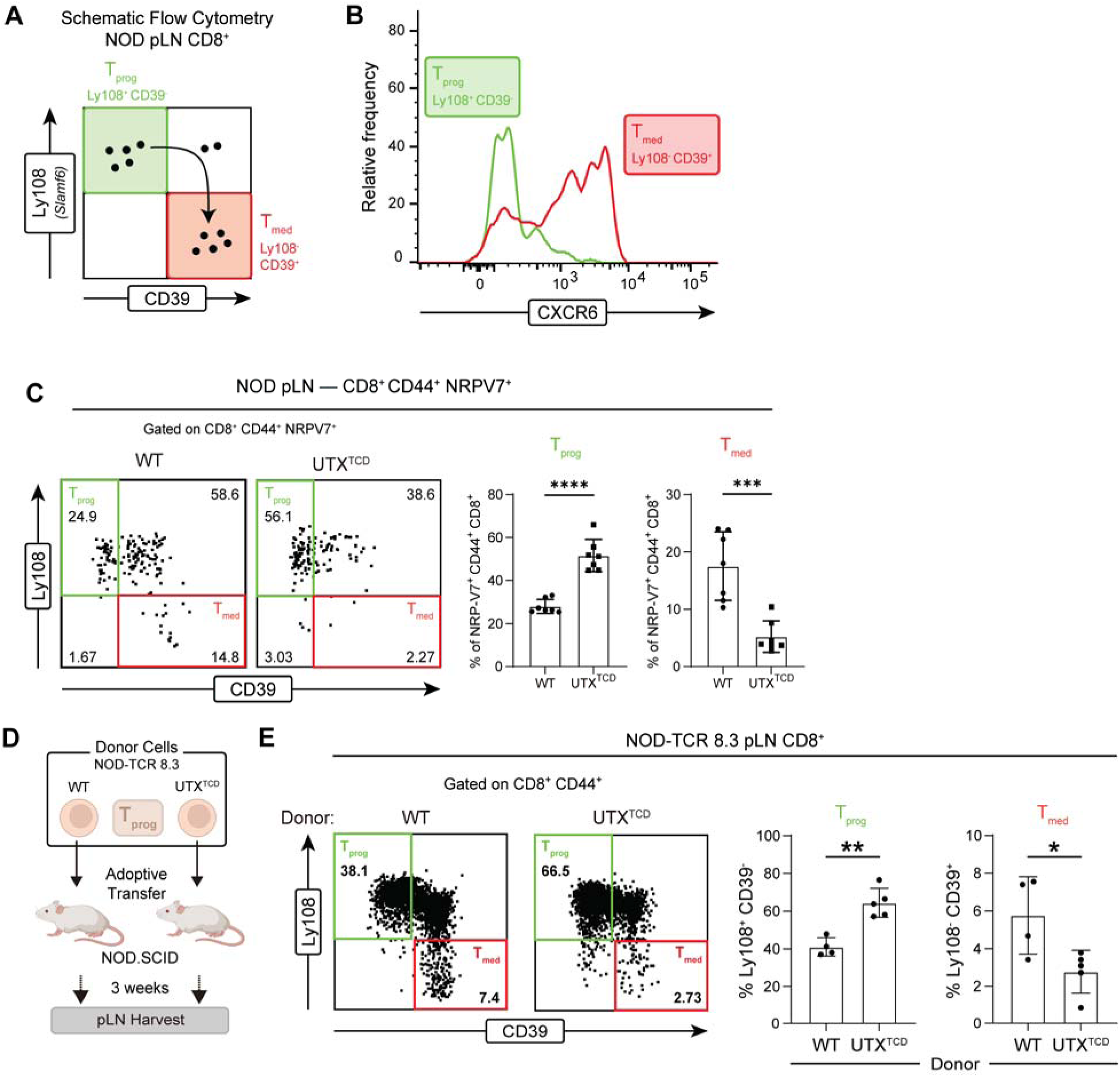
UTX promotes T_prog_ to T_med_ conversion of beta cell-specific CD8^+^ T cells. **(A)** Gating scheme for identifying T_prog_ and T_med_ cells by flow cytometric analysis. **(B)** Representative histogram of CXCR6 expression in T_prog_ vs. T_med_ cell populations identified by Ly108 and CD39 expression. (**C)** Representative flow cytometric plot (left), and average frequencies (right) of T_prog_ (Ly108^+^ CD39^-^) and T_med_ (Ly108^-^ CD39^+^) cells within antigen-experienced, IGRP-specific pLN CD8^+^ T cells (CD8^+^ CD44^+^ NRP-V7^+^) of *NOD*-*WT* and *NOD-UTX*^TCD^ female littermates (15-29 weeks of age). ****p<0.0001; ***p<0.001; Student’s t test. (**D)** Scheme for adoptive transfer of pLN T_prog_ cells (Ly108^+^ CD39^-^) from *NOD-TCR 8.3 WT* vs. *NOD-TCR 8.3 UTX^TCD^* mice into *NOD.SCID* recipients. (**E)** Representative flow cytometric plot (left) and average frequencies (right) of T_prog_ [Ly108^+^ CD39^-^] or T_med_ [Ly108^-^ CD39^+^] among pLN antigen-experienced (CD8^+^ CD44^+^) cells in hosts after adoptive transfer as shown in (D). **p<0.01; *p<0.05; Student’s t test.

To further delineate UTX’s functions within beta cell-specific T_prog_ cells, we performed adoptive transfers of IGRP-specific T_prog_ into immunodeficient *NOD.SCID* recipients (**Figure 3D**). To isolate sufficient numbers of IGRP-specific T_prog_ for transfer, we exploited an NOD T cell receptor (TCR) transgenic mouse line (*NOD-TCR 8.3*) in which T cells express a TCR recognizing the IGRP_206-214_ epitope in the context of MHC Class I (24). IGRP-specific CD8+ T cell progenitors (Ly108^+^ CD39^-^) were sorted from 6-week-old *NOD-TCR 8.3* mice with or without T cell-specific deficiency in UTX (*NOD-TCR 8.3 UTX^TCD^ or NOD-TCR 8.3 WT*, respectively). Three weeks after transfer, an increased frequency of pLN T_prog_ cells (Ly108^+^ CD39^-^ among CD8^+^ CD44^+^) was seen in hosts receiving UTX-deficient CD8^+^ T_prog_ cells, compared to wildtype (**Figure 3E**). Concomitantly, a decrease in the frequency of pLN T_med_ (Ly108^-^ CD39^+^ among CD8^+^ CD44^+^) was noted in recipients of UTX-deficient T_prog_ cells (**Figure 3E**). Together, these findings support a model in which UTX functions in IGRP-specific CD8^+^ T_prog_ cells to promote their transition to T_med_ cells.

We next asked whether UTX may play a parallel function in CD8^+^ T_prog_ cells within the pancreas. To address this, we performed flow cytometric analysis of pancreatic CD8^+^ T cell subsets in *NOD-TCR 8.3 UTX^TCD^* vs. *NOD-TCR 8.3 WT* mice **(Supplemental Figure 5A).** Similar to pLN CD8^+^ T cells, pancreatic CD8^+^ T cells showed altered subset distribution with UTX deficiency. T_prog_ cells (Ly108^+^ CD39^-^) were increased, while T_med_ cells (Ly108^-^ CD39^+^) were decreased, in the pancreas of *NOD-TCR 8.3 UTX^TCD^*mice compared to *NOD-TCR 8.3 WT* mice. These findings suggest that UTX’s role in CD8^+^ T_prog_ cells in the pancreas mirrors its role in the pLN.

### UTX controls chromatin accessibility and transcriptional changes in antigen-specific pLN T_prog_ cells

Recent studies have demonstrated the importance of epigenetic and transcriptional regulatory programs in controlling the transition of CD8^+^ T cell states (25). Because UTX is an epigenetic regulator (26), we reasoned that UTX may control T_prog_ to T_med_ conversion by remodeling chromatin at regulatory elements of target gene loci. To examine this, we performed bulk ATAC-seq and RNA-seq in parallel on sort-purified beta cell-specific CD8^+^ T_prog_ cells from pLN of either *NOD-TCR 8.3 UTX^TCD^* or *NOD-TCR 8.3 WT* mice. ATAC-seq analysis demonstrated large-scale chromatin accessibility alterations in UTX-deficient CD8^+^ T cells. Principal component analysis (PCA) plots showed sample clustering by genotype (**Figure 4A, top),** highlighting the extensive impact of UTX on the chromatin landscape of CD8^+^ T cells. RNA-seq analysis performed in parallel also showed large-scale transcriptional changes, with PCA analysis of gene expression demonstrating genotype-specific clustering (**Figure 4A, bottom)**. ATAC-seq revealed 4,075 peaks decreased in accessibility and 8,807 peaks increased in accessibility in *NOD-TCR 8.3 UTX^TCD^* progenitors, compared to UTX-sufficient controls (|log_2_ fold change|L>L0.5, false discovery rate (FDR)-adjusted PL<L0.05) **(Data File S2)**. These differentially accessible peaks corresponded to 6,859 differentially accessible genes. Further, RNA-seq revealed 1,280 decreased and 1,768 increased transcripts in *NOD-TCR 8.3 UTX^TCD^* progenitors, compared to UTX-sufficient controls (|log_2_ fold change|L>L0.5, FDR-adjusted PL<L0.05) **(Data File S3)**.

**Figure 4.**
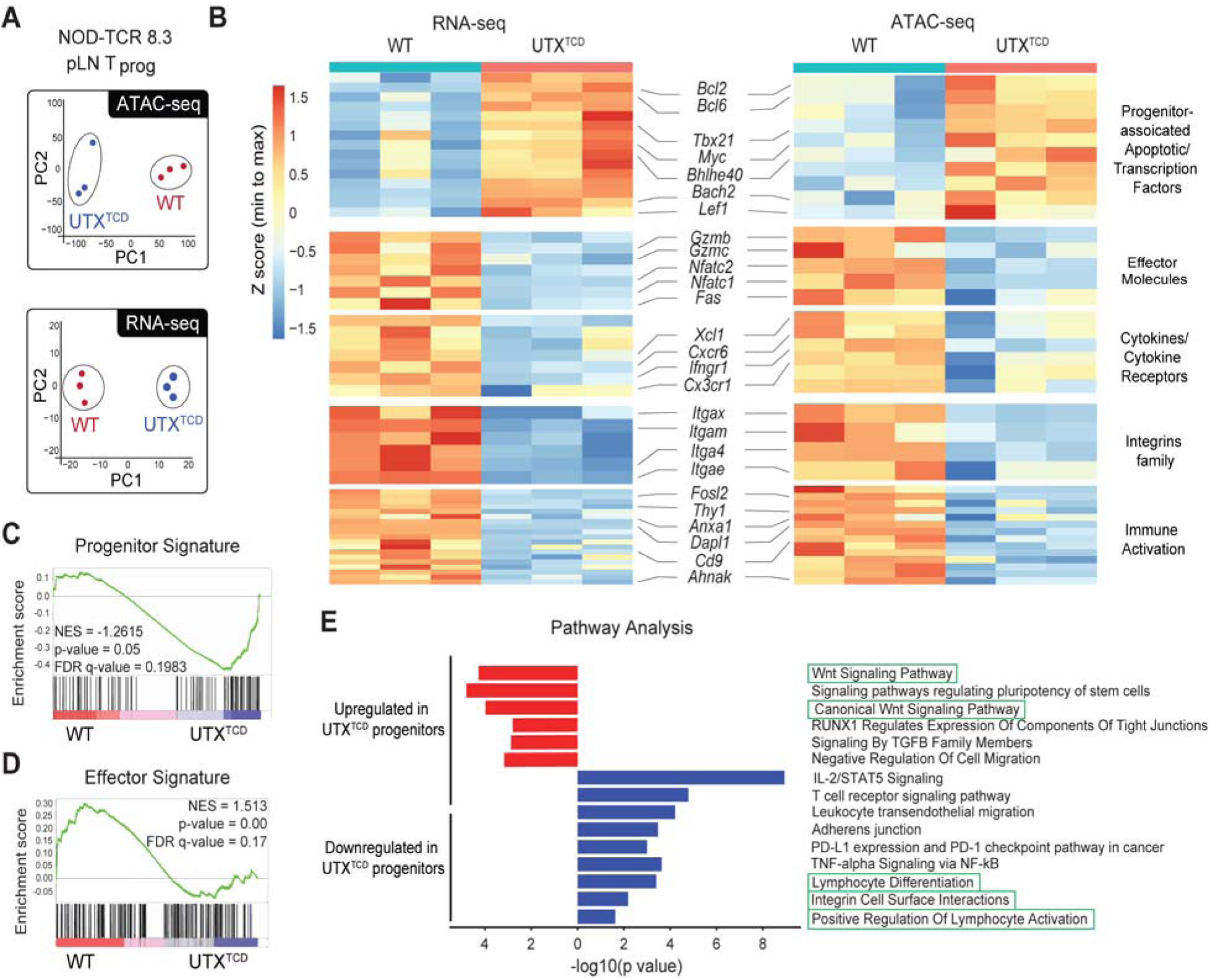
UTX controls chromatin accessibility at progenitor and effector gene loci in antigen-specific pLN T_prog_ cells. **(A)** PCA analysis of ATAC-seq (top) and RNA-seq (bottom) profiles of sorted T_prog_ (CD8^+^ Ly108^+^ CD39^-^) from pLN of *NOD-TCR 8.3 WT* and *NOD-TCR 8.3 UTX^TCD^* (n = 3/group). **(B)** Heatmaps of the expression (left, RNA-seq) and chromatin accessibility (right, ATAC-seq) of selected differentially expressed and accessible genes involved in immune function. The color scale indicates the relative expression or accessibility, with red representing higher values and blue representing lower values. (**C and D)** Gene set enrichment analysis (GSEA) for Progenitor signature (**Data File S4**) (C) and Effector signature (D) genes in the RNA-seq dataset of T_prog_ from *NOD-TCR 8.3 WT* and *NOD-TCR 8.3* UTX^TCD^. **(E)** Pathway analysis of differentially accessible genes by ATAC-seq using Enrichr (p-value by Fisher’s exact test).

Integrative analysis of ATAC- and RNA-seq identified a significant positive correlation (Pearson R = 0.73, p < 2.2×10^−16^) between genotype-dependent changes in chromatin accessibility and gene expression **(Supplemental Figure 6A)**. These data suggest a direct correlation between UTX-mediated alterations in chromatin accessibility and transcription. Concurrent changes in chromatin accessibility and transcription were observed in genes with known roles in the differentiation and maintenance of stem-like progenitor CD8^+^ T cells. For instance, *NOD-TCR 8.3 UTX^TCD^* T_prog_ cells demonstrated increased chromatin accessibility, with corresponding increased gene expression, at the *Bcl6* locus (**Figure 4B and Supplemental Figure 6B)**, a transcription factor that prolongs stem-like progenitor persistence (27). Additionally, *NOD-TCR 8.3 UTX^TCD^*T_prog_ also showed increased chromatin accessibility and gene expression at the *Bach2* locus (**Figure 4B and Supplemental Figure 6B)**, a transcriptional repressor that promotes the stem-like CD8^+^ T cell program, while suppressing terminal exhaustion programs (27). Finally, *Bcl2*, an anti-apoptotic factor that promotes stem-like progenitor cell survival (28), also showed increased accessibility and expression in *NOD-TCR 8.3 UTX^TCD^* T_prog_ cells (**Figure 4B and Supplemental Figure 6B)**. Thus, UTX deficiency is associated with changes at loci that are important in stem-like progenitor maintenance and differentiation.

At the same time, decreased accessibility and reduced expression were seen in *NOD-TCR 8.3 UTX^TCD^* T_prog_ cells at genes important for CD8^+^ T cell effector function (**Figure 4B and Supplemental Figure 6B)**. These genes include cytolytic molecules (e.g., *Gzmb, Gzmc, Fas*) important in beta cell killing (29); integrins (e.g., *Itga4*, *Itgae*) (30, 31) critical in leukocyte islet infiltration; and cytokine/cytokine receptors (e.g., *Ifngr1, CXCR6*) (4, 32, 33). Both *Dapl1* and *Nfatc2,* which function together in CD8^+^ T cell effector function (34), were also less accessible and expressed. Moreover, genes associated with T cell activation (*Fosl2, Thy1, Cd9*) were also less accessible and downregulated in expression in CD8^+^ T cells from *NOD-TCR 8.3 UTX^TCD^* mice (35–37). Gene set enrichment analysis (GSEA) confirmed these findings, showing upregulated stem-like ‘Progenitor’ and downregulated ‘Effector’ gene signatures in *NOD-TCR 8.3 UTX^TCD^* cells (**Figure 4, C and D and Data File S4)** (4, 9).

Enrichment analysis of differentially accessible genes revealed association with pathways important in stem-like progenitor generation as well as cytolytic effector differentiation and function. For instance, Wnt signaling is critical in the generation and maintenance of stem-like memory T cells (38), and Enrichr pathway analysis of differentially accessible genes revealed upregulation of ‘Wnt Signaling Pathway’ and “Canonical Wnt Signaling Pathway’ genes in *NOD-TCR 8.3 UTX^TCD^*T_prog_ cells (**Figure 4E**). At the same time, ‘Lymphocyte Differentiation’, ‘Integrin Cell Surface Interactions’, ‘Positive Regulation of Lymphocyte Activation’, and other pathways associated with cytolytic effector function were downregulated in *NOD-TCR 8.3 UTX^TCD^* T_prog_ cells (**Figure 4E**). Together, these findings provide further support that UTX poises chromatin in stem-like T_prog_ cells for transition to cytolytic effectors.

### Integrated analysis of UTX chromatin binding, accessibility, and transcription reveals STAT3 and TCF1 co-occupancy at UTX target genes

The genome-wide changes in accessibility and transcription in UTX-deficient T_prog_ cells may be attributed either to i) UTX’s direct regulation of chromatin accessibility and gene expression, or ii) secondary changes that result from alterations in other UTX-mediated epigenetic and transcriptional regulators. To identify gene loci directly regulated by UTX, we utilized UTX CUT&Tag (Cleavage Under Targets and Tagmentation Assay) (39) to identify gene loci bound by UTX on sort-purified *NOD-TCR 8.3 WT* T_prog_ cells. Sort-purified *NOD-TCR 8.3 UTX^TCD^* T_prog_ cells were used as a negative control. PCA of UTX CUT&Tag data revealed sample clustering by genotype (**Figure 5A**). We identified 6,718 UTX-bound peaks (FDR-adjusted P < 0.05), which corresponded to 5,222 distinct genes **(Data File S5)**. Genomic annotation of UTX CUT&Tag peaks demonstrated a strong promoter bias, with 64.5% of UTX-bound regions localized within 1□kb of transcription start sites (**Figure 5B**). This promoter-proximal enrichment suggests that UTX primarily engages core regulatory elements to exert its epigenetic functions in T_prog_ cells.

**Figure 5.**
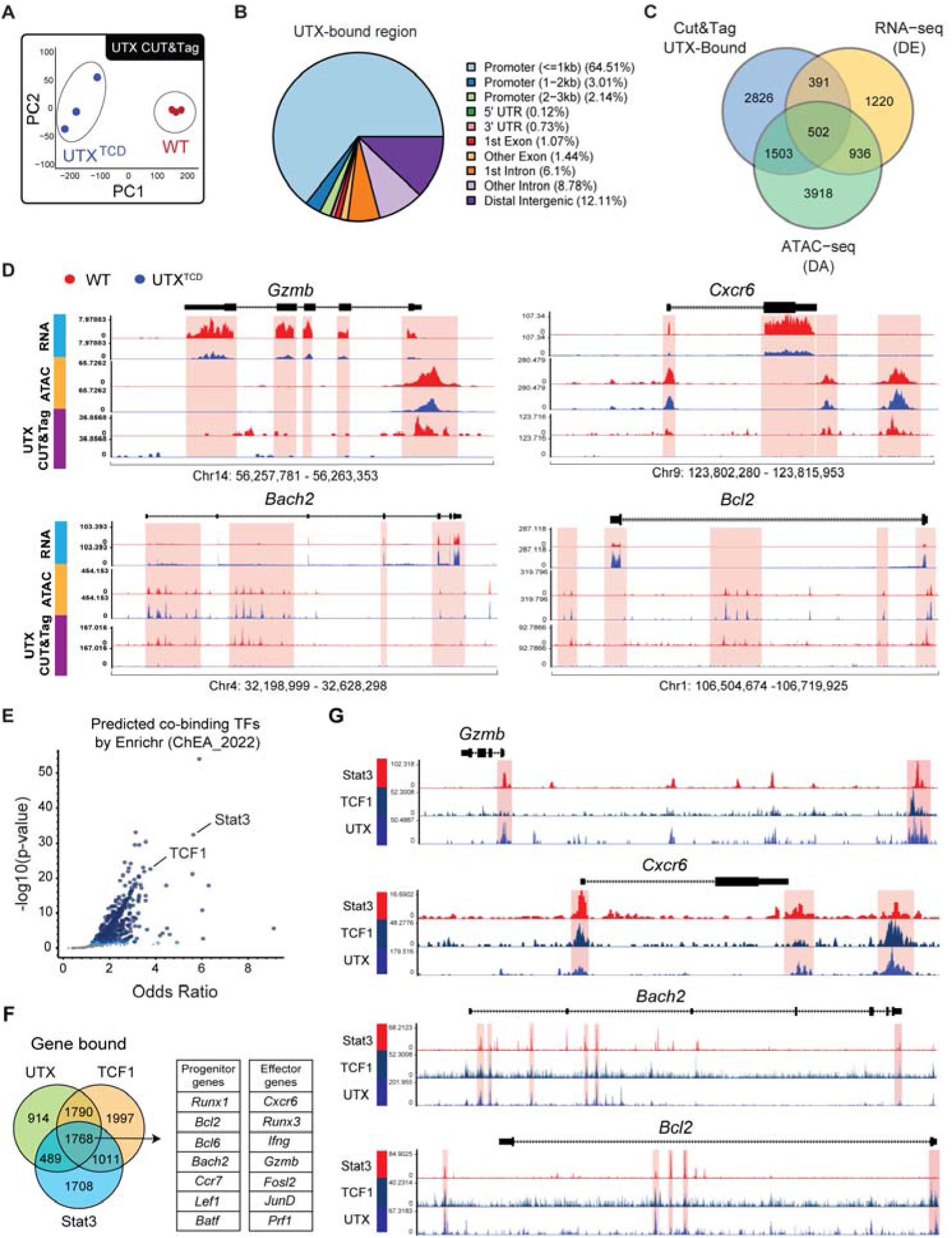
Integrative analysis of chromatin binding, accessibility, and transcription reveals STAT3 and TCF1 co-bind UTX-target genes. **(A)** PCA analysis of UTX CUT&Tag from sorted T_prog_ (CD8^+^ Ly108^+^ CD39^-^) from pLN of *NOD-TCR 8.3 WT* and *NOD-TCR 8.3 UTX^TCD^*. **(B)** Location of UTX-bound peaks. **(C)** Venn diagram outlining overlapping differentially accessible (DA) genes identified by ATAC-seq and differentially expressed (DE) genes identified by RNA-seq. **(D)** Representative gene tracks from the UCSC Integrated Genome Browser of anti-UTX CUT&Tag, ATAC-seq, and RNA-seq of effector genes, *Gzmb* and *Cxcr6*, and progenitor genes, *Bach2* and *Bcl2*. The Y-axis depicts counts per million (CPM). **(E)** Transcriptional factors were bound to the overlapping genes identified by UTX datasets. Overlapping genes were input into Enrichr for ChEA3 analysis. **(F)** Venn diagram outlining overlapping genes bound by UTX, TCF1, and STAT3. (TCF1-ChIP-seq: GSE73240, STAT3-ChIP-seq: GSE217374). **(G)** Gene tracks from the UCSC genome browser to demonstrate the aligned region bound by UTX, TCF1, and STAT3. The Y-axis depicts counts per million (CPM).

We utilized this UTX CUT&Tag dataset, the ATAC-seq dataset (**Figure 4B**), and the RNA-seq dataset (**Figure 4B**) to identify 502 gene loci that are UTX-bound, differentially accessible, and differentially expressed in UTX-deficient T_prog_ cells (**Figure 5C**). These UTX-bound genes included stem-like progenitor-associated genes (i.e., *Bach2, Bcl2, Bcl6*) that are increased in accessibility and transcription within *NOD-UTX^TCD^*T_prog_ cells (**Figure 5D and Supplemental Figure 7A)**, suggesting that UTX binds directly to stem-like progenitor-associated genes to decrease accessibility and repress transcription at these loci. Additionally, UTX-bound genes included mediator-associated genes (i.e., *Gzmb, Cxcr6, Thy1*) that are decreased in accessibility and transcription within *NOD-UTX^TCD^* T_prog_ cells (**Figure 5D and Supplemental Figure 7A)**. These analyses demonstrate that in antigen-specific T_prog_ cells, UTX poises chromatin to repress transcription at key T_prog_ genes and to promote transcription at effector genes.

While UTX has intrinsic histone demethylase activity, our data demonstrate that UTX’s demethylase activity is dispensable in T1D pathogenesis **(Supplemental Figure 1B)**. An alternative mechanism by which UTX exerts its regulatory effects is through interactions with transcription factors (TFs) (40). To identify TF binding partners of UTX, we used a TF enrichment analysis tool (ChEA3 in Enrichr) to prioritize TFs potentially responsible for the observed 502 UTX-bound, differentially accessible, and differentially expressed target genes (**Figure 5E**) (41). Highly associated TFs included i) TCF1, a key regulator of stem-like progenitors (9) and ii) STAT3, a TF implicated in T_prog_ to effector conversion and a key TF in T1D pathogenesis (10–12). These findings suggest the possibility that UTX may function in concert with TCF1 and STAT3 to regulate differentiation of T_prog_ cells to an effector state.

We next integrated our UTX CUT&Tag dataset with publicly available chromatin immunoprecipitation datasets to determine whether gene loci may be co-occupied by UTX, TCF1, and STAT3 proteins. Cross-referencing of UTX-bound genes with TCF1-bound (GSE73240) and STAT3-bound (GSE217374) genes showed substantial overlap of 1,768 genes (**Figure 5F**). These overlapping genes included progenitor-associated genes (*Bcl2, Bach2, Bcl6, and Lef1*) and mediator-associated genes (*Cxcr6, Gzmb, Ifngr1, and Thy1*) that are differentially accessible and differentially expressed in *NOD-UTX^TCD^*CD8^+^ T_prog_ cells, compared to *NOD-WT*. Moreover, comparison of UTX-bound, TCF1-bound, and STAT3-bound peaks showed a high degree of overlap in multiple genes (**Figure 5G and Supplemental Figure 8A).** Binding by UTX, TCF1, and STAT3 was noted at key effector-associated genes (e.g., *Gzmb*, *Cxcr6*, *Runx3*, and *Ifng*) as well as progenitor-associated genes (e.g., *Bach2*, *Ccr7*, and *Lef1)*, suggesting co-occupancy of UTX, TCF1, and STAT3 at these target gene loci. Collectively, these data suggest that UTX cooperates with TCF1 and STAT3 to influence the transition of antigen-specific T_prog_ to an effector state.

### UTX promotes differentiation of human stem-cell memory CD8^+^ T cells into a terminally differentiated effector state through binding with TCF1 and STAT3

Recent work has associated T1D in humans with increased circulating CD8^+^ T memory stem cells (T_SCM_), a long-lived, self-renewing beta cell-specific population (2, 42, 43). T_SCM_ cells play a key role in T1D pathogenesis by giving rise to terminally differentiated effector/mediator (T_TE_) cells that ultimately kill insulin-producing beta cells. Thus, the T_SCM_ (CCR7^+^ CD45RA^+^ CD95^+^) population in T1D patients may have a parallel role to the T_prog_ population in NOD mice, while the T_TE_ (CCR7^-^ CD45RA^+^ CD95^+^) population in T1D patients may have a parallel role to the T_med_ population in NOD mice (42, 43).

To test whether UTX controls T_SCM_ to T_TE_ conversion, we utilized CRISPR/Cas9 to edit the *UTX* locus in CD8^+^ T cells isolated from T1D patient peripheral blood mononuclear cells. We confirmed efficient knockdown of UTX protein expression using UTX-specific sgRNAs compared to a non-targeting control (NTC sgRNA) (**Figure 6A**). Following a seven-day culture with homeostatic cytokines (IL-2, IL-7, IL-15), we observed an increased frequency of T_SCM_ cells (CCR7^+^ CD45RA^+^) and a corresponding decrease in T_TE_ cells (CCR7^-^ CD45RA^+^) in UTX-deficient CD8^+^ T cells relative to controls (**Figure 6B**). This altered distribution supports a cell-intrinsic role for UTX in governing T_SCM_ to T_TE_ conversion in human CD8^+^ T cells, indicating a conserved, cross-species function for UTX.

**Figure 6.**
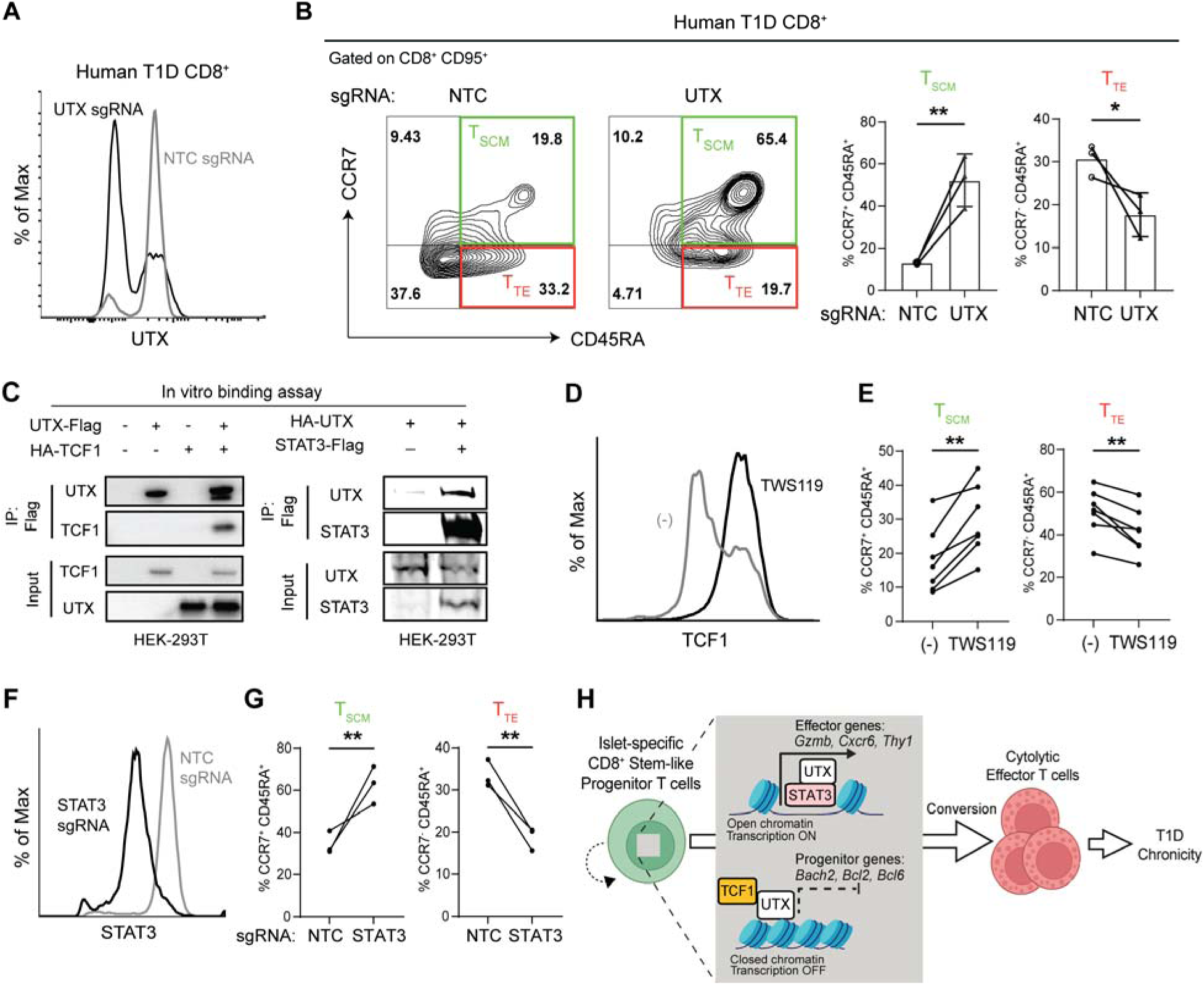
UTX regulates human stem-cell memory CD8^+^ T cell differentiation into the terminally effector state and interacts with TCF1 and STAT3. **(A)** Representative flow cytometric plot of CD8^+^ T cells from patients with T1D after CRISPR-Cas9 editing with NTC (non-targeting control) sgRNA or UTX-targeting sgRNA. **(B)** Representative flow cytometric plots (left) and average frequency (right) of T_SCM_ and T_TE_ cells among CD8^+^ CD95^+^ peripheral blood cells from T1D patients. Cells were edited with either NTC sgRNA or UTX-targeting sgRNA. **(C)** In vitro binding assay to show the interaction of UTX-TCF1 (left) and UTX-STAT3 (right) in HEK-293T cells by co-transfection and co-immunoprecipitation. **(D)** Representative flow cytometric histogram plot of TCF1 expression in CD8^+^ T cells treated with DMSO or 5uM of TWS119 supplemented with IL-7 (10ng/mL) and IL-21 (25ng/mL) for 7 days. **(E)** Frequencies of T_SCM_ and T_TE_ within CD8^+^ CD95^+^ T cells treated with DMSO or 5uM of TWS119 for 3 days. Each dot represents a biological replicate (n=7). **p<0.01; paired Student’s t test. **(F)** Representative flow cytometric histogram plot of CD8^+^ T cells from patients with T1D after CRISPR-Cas9 editing with NTC (non-targeting control) sgRNA or STAT3-targeting sgRNA. **(G)** Frequencies of T_SCM_ and T_TE_ within CD8^+^ CD95^+^ T cells after CRISPR/Cas9 editing with sgSTAT3 versus a non-targeting control (NTC). Left panel: T_SCM_; right panel: T_TE_. Each dot represents a biological replicate (n=3). **p<0.01; paired Student’s t test. **(H)** Model of how UTX forms a complex with TCF1 and STAT3 to modulate the progenitor and effector programs in promoting T1D development.

Given that our integrative analysis suggested a high degree of cooperativity between UTX, TCF1, and STAT3 (**Figure 5, E-G)**, we utilized a co-immunoprecipitation approach to assess for binding of UTX to TCF1 and STAT3. HEK-293T cells were co-transfected with epitope-tagged constructs encoding UTX in combination with either TCF1 or STAT3, and subsequent immunoprecipitation demonstrated interactions between UTX and both TCF1 and STAT3 (**Figure 6C**). In line with this finding, endogenous UTX protein co-immunoprecipitated with TCF1 and STAT3 in Jurkat cells **(Supplemental Figure 9A).** These data demonstrate that UTX directly binds critical transcription factors important in progenitor specification (9) and progenitor-to-effector differentiation (10–12).

TCF1 expression is induced by inhibitors of glycogen synthase kinase 3 beta, such as TWS119 (9). Consistent with TCF1’s role in enforcing progenitor formation and maintenance, treatment of T1D patients’ CD8^+^ T cells with TWS119 resulted in the induction of TCF1 (**Figure 6D**) and an increased frequency of T_SCM_ and a decreased frequency of T_TE_ cells (**Figure 6E**). Moreover, STAT3 drives progenitor T cell conversion to effectors (11), and editing of the STAT3 locus in T1D patients’ CD8^+^ T cells resulted in accumulation of T_SCM_ cells and decreased T_TE_ cells (**Figure 6, F and G).** Based on these data, we propose a mechanistic model wherein UTX interacts with STAT3 to enhance effector gene transcription necessary for the progenitor-to-effector transition, while simultaneously binding TCF1 to suppress its progenitor-maintaining functions during differentiation (**Figure 6H**). These factors function in tandem at gene loci important in effector function and progenitor identity, insight that enhances our understanding of the molecular mechanisms underpinning autoimmune chronicity and offers potential avenues for targeted therapeutic interventions in T1D.

## DISCUSSION

For patients with T1D, the persistence of the beta cell autoimmune response results in a lifetime of insulin dependence. As a consequence, patients are predisposed to cardiovascular disease and other T1D-associated complications, including shortened life expectancy (44). Accumulating evidence points to a key role for stem-like T_prog_ cells in prolonging the autoimmune response (2, 3). These long-lived T_prog_ cells continually repopulate the pool of terminally differentiated, cytolytic CD8^+^ T cells that destroy beta cells. The mechanisms controlling the conversion of T_prog_ cells to cytolytic effectors, however, remain incompletely understood. Here, we identify UTX as an epigenetic regulator that primes T_prog_ cells for transition to cytolytic effectors. UTX exhibits regulatory function in CD8^+^ T cell differentiation through its interactions with TCF1 and STAT3 transcription factors, independent of its intrinsic histone demethylase activity. Taken together, these findings suggest that this UTX:TCF1:STAT3 regulatory complex may be targeted therapeutically to disrupt the chronic autoimmune process of T1D.

In beta cell-specific T_prog_ cells, UTX represses transcription at gene loci (*Bcl6, Bach2, Bcl2*) well-described for their roles in the differentiation and maintenance of stem-like progenitors. At the same time, UTX organizes chromatin for enhanced transcription at multiple effector loci (e.g., *Gzmb, Thy1, Cxcr6*) in CD8^+^ T_prog_ cells. These data point to UTX’s dual role of repressing stem-like progenitor maintenance in favor of promoting a cytolytic effector CD8^+^ T cell program. Our findings suggest that coordination with TCF1 and STAT3, transcription factors with respective functions in progenitor specification (9) and progenitor-to-effector differentiation (10–12), is critical for the function of UTX in diabetogenic CD8^+^ T cells.

Genetic association studies in T1D have been instrumental in identifying pathogenic mechanisms. The Type 1 Diabetes Genetics Consortium (T1DGC) reported that a *TCF7* C883A polymorphism was associated with T1D in patients who do not carry the high-risk HLADR3/4 genotype (7). This finding corroborated a previous report from the T1DGC Rapid Response Project that linked T1D with the *TCF7* locus (8). In addition to *TCF7*, alterations in *STAT3* also have a notable impact on T1D susceptibility (45). Approximately 30% of patients with gain-of-function *STAT3* mutations develop T1D (46), and NOD mice with a human gain-of-function *STAT3* mutation develop accelerated T1D (10, 12). Indeed, our findings indicate that UTX forms a complex with TCF1 (encoded by *TCF7*) and STAT3 to promote the differentiation of progenitor CD8^+^ T cells into an effector state, contributing to the autoimmune response in T1D.

How STAT3 contributes to T1D pathogenesis has been the subject of much interest. Early studies suggested that STAT3 gain-of-function mutations result in changes to the distribution of CD4^+^ T cell subsets, noting increased ratios of T helper 17 (Th17) to regulatory T cells (Treg) (46). In contrast, recent studies have demonstrated alterations in the CD8^+^ T cell compartment, such that effector CD8^+^ T cells were increased in mice harboring human STAT3 gain-of-function mutations (10, 12). While possible that STAT3-mediated changes to both CD4^+^ and CD8^+^ T cells may underlie the promotion of T1D, our findings support the important role of STAT3 in CD8^+^ T cell differentiation by coordination with UTX and TCF1.

While our data demonstrate that UTX promotes the diabetogenic capacity of CD8^+^ T cells and the conversion of CD8^+^ T_prog_ to T_med_ cells, we cannot exclude the possibility that UTX also plays a role in CD4^+^ T cells during T1D pathogenesis. Indeed, UTX has previously been shown to upregulate IL-6R expression in CD4^+^ T cells, promoting IL-6-dependent T follicular helper (Tfh) differentiation in the context of chronic LCMV (lymphocytic choriomeningitis virus) infection (18). However, we did not see a difference in the frequency of CD4^+^ Tfh cells in the pLN of *NOD-WT* vs. *NOD-UTX^TCD^* mice. We speculate that this reflects a difference in the dependence of these two model systems on IL-6. While IL-6 is required for late-stage control of chronic LCMV infection (55), IL-6 deficiency does not impair diabetes development in NOD mice (56). This suggests that other cytokines besides IL-6 may play a role in the differentiation of Tfh cells in NOD mice. Indeed, IL-21 has been identified as a redundant cytokine to IL-6 in the induction of Tfh cells (57). Additionally, CD4^+^ T cells with stem-like features that express TCF1 have recently been described in NOD mice (19), and whether UTX may also regulate this CD4^+^ T cell subset requires further investigation.

Between 2002-2012, the incidence of T1D in youth increased by 1.4% per year (47), a rise that prompts us to question what factors may be driving the development of autoimmune disease beyond genetic susceptibility. Epigenetic alterations have been indicated as the possible bridge between environmental influence and genetics in the development of disease, such that environmental cues induce alterations to gene expression without changes to the DNA sequence itself (48). In this study, we show that diabetogenic CD8^+^ T cells and their developmental trajectory are orchestrated by the epigenetic regulator UTX. Our findings suggest that therapies blocking cytolytic effector CD8^+^ T cell differentiation and function may be effective therapeutic strategies for T1D. Therapies that modulate epigenetic mechanisms are already in use for cancer (49), and our data reveal the UTX:TCF1:STAT3 complex as a potential therapeutic target for chronic autoimmunity.

## MATERIALS AND METHODS

### Sex as a biological variable

Only females were included in these studies since UTX is X-linked and expression of the Y-linked homolog UTY in males could confound analyses.

### Mice

*NOD/ShiLtJ* (NOD; JAX:001976) and *NOD.Cg-Prkdc^scid^/J* (*NOD.SCID*; JAX:001303) were purchased from The Jackson Laboratory. *NOD-UTX^fl/fl^.LckCre+* (*NOD-UTX^TCD^*) was generated by backcrossing *B6-UTX^fl/fl^ LckCre+* mice to NOD mice > 10 generations (50). We confirmed that the 26 identified diabetes associated (Idd) loci were derived from recipient NOD strain by congenic fine mapping (MegaMUGA). Cre-negative littermates were used as UTX-sufficient, wildtype controls (*NOD-WT*). *NOD-TCR 8.3 UTX^TCD^* were generated by crossing *NOD-UTX^TCD^*mice with *NOD.Cg-Tg (TcraTcrbNY8.3)1Pesa/DvsJ* (*NOD-TCR 8.3*; JAX:005868). All mice were housed in a specific pathogen-free barrier facility at UCLA. Experiments were performed in compliance with UCLA Animal Research Committee regulations.

The *NOD.UTX^DMD^* mice were generated at the UNC Animal Models Core. These mice have mutations that result in amino acid alterations at H1146A and E1148A that inactivate the demethylase activity of UTX (54). These mutations were achieved by injecting NOD pro-nuclei with gRNA:Cas9 complexes that target introns flanking exon24, which encodes the JmjC domain, along with a plasmid donor encoding the mutated exon24. Single-cell mouse embryos were then surgically implanted into the oviducts of pseudopregnant recipient NOD mice. The resulting pups were further backcrossed to NOD prior to intercrossing to generate female homozygous mice for use in these studies. Mice with the knock-in allele were identified by PCR and sequencing using PCR primers: (Utx-Intron23-ScF2: CACGCATTTCCCAGC ACTTG; Utx-Int23-ScR2: CCTACTTTTCACAGAAGTCATTCAAAACAC), and sequencing using primer: (Utx-Int23-SqF1: TAACAAATATATCTTATTGGGCACC). These mice were housed in specific pathogen-free facilities at UNC-Chapel Hill, and experiments performed with them were approved by the Institutional Animal Care and Use Committee at UNC-Chapel Hill.

### Assessment of diabetes

Mice were monitored for diabetes at least once per week using urine glucose strips (Ascensia Diabetes Care). Mice were considered diabetic after two consecutive urine tests matched with the color of glucose concentration ≥ 250 mg/l.

### Adoptive T cell transfer

All adoptive transfer experiments used *NOD.SCID* mice as hosts and were performed by intravenous tail vein injections. For CD8^+^ T cell transfers, splenic CD8^+^ T cells of *NOD-WT* or *NOD-UTX^TCD^*mice were isolated by EasySep™ Mouse CD8^+^ T Cell Isolation Kit (STEMCELL Technologies, #19853), and splenic CD4^+^ T cells from *NOD-WT* mice were isolated by EasySep™ Mouse CD4^+^ T Cell Isolation Kit (STEMCELL Technologies, #19852). 1×10^6^ splenic CD8+ T cells were co-transferred with 1×10^6^ splenic CD4^+^ T cells into the host. Diabetes development was assessed twice a week.

For CD8^+^ T_prog_ transfer, Ly108^+^ CD39^-^ among CD8^+^ CD44^+^ T_prog_ were sorted from the pLN of *NOD-TCR 8.3 UTX^TCD^* or *NOD-TCR 8.3 WT* mice. 250,000 CD8^+^ T_prog_ were co-transferred with 1×10^6^ splenic WT CD4^+^ T cells into the hosts. Three weeks post-transfer, pLN CD8^+^ T cells were analyzed by flow cytometry.

### Histology and insulitis scoring

Hematoxylin and eosin (H&E) staining of pancreas tissues are done by UCLA Translational Pathology Core Laboratory (TPCL). Insulitis for each pancreatic islet is scored based on percent of islet infiltrated by immune cells: 0 = no infiltration, 1 = 0 – 25% infiltration, 2 = 25 – 50% infiltration, 3 = 50 – 75% infiltration, 4 = 75 – 100% infiltration.

### Immune cell isolation

Mice were euthanized by CO_2_. Lymph nodes and spleens were mechanically disrupted with the back of a 1 mL syringe, filtered through a 40-µm strainer, and subjected to ACK lysis for 1 min. Cells were washed once with PBS. Pancreas digestion was adapted from existing protocols. In brief, 3 mL of collagenase type IV solution (2mg/ml) with DNaseI (10U/ml) in HBSS (Gibco 14025-092) supplemented with 10% fetal bovine serum (FBS) and DNaseI (10U/ml) was injected into the clamped pancreatic duct. Perfused pancreases were then excised and incubated in 4 mL of the collagenase solution at 37°C for 30 min. The digested pancreas was then washed using 10mL HBSS with 10% FBS and spun down at 300g for 3 min, 2 times, followed by mechanically breaking up the pancreas. Then, tissue was passed through a 40-µm strainer and washed with HBSS with 10% FBS, and spun down at 400g for 5 min. Finally, the pellet was resuspended in FACS buffer for antibody staining.

### Flow cytometry and cell sorting

Cells were analyzed for surface markers using fluorophore-conjugated antibodies. Cell surface staining was performed in FACS buffer (2% FBS and 2mM EDTA in PBS) for 30 min at 4°C. Intranuclear protein staining, including UTX, H3K27me3, and TCF1, was performed by fixing and permeabilizing using the eBioscience Foxp3/Transcription Factor kit. Followed by the primary rabbit antibody, Goat Anti-Rabbit IgG H&L (Abcam) was used as the secondary antibody.

H-2Kd/NRP-V7 (KYNKANFL) tetramer was obtained from the NIH Tetramer Core Facility. Tetramer staining was performed with two steps: cells were stained with tetramer in PBS solution (1:200) for 30 min at room temperature, followed by staining with surface markers for 30 min at 4°C. For intracellular cytokine staining, cells were stimulated with PMA (50ng/mL), ionomycin (1μg/mL), Brefeldin A (1×), and monensin (1×) for 4 h. Cells were stained with surface antigen-targeting antibodies, then fixed and permeabilized using the BD Cytofix/Cytoperm Fixation/Permeabilization Kit (BD Biosciences, 554714) for subsequent cytokine staining. Flow cytometry was performed on an LSRII Fortessa, and FACS sorting was performed on an ARIA or ARIA-H instrument (BD Biosciences, San Jose, CA) at the UCLA Broad Stem Cell Research Center Flow Cytometry Core. The data were analyzed with FlowJo v10.7.2 (TreeStar). Antibodies used are shown in **Table S1**.

### Single cell RNA analysis

Pancreatic lymph node (pLN) scRNAseq data are clustered using *Seurat* package (version 4.4.0). Differential gene markers of each cluster are computed using the FindAllMarkers function of Seurat. Immune cell types are defined based on their canonical markers (**Data File S1A**). Cd8 T cell clusters, including clusters with mixed Cd8 and Cd4 T cells, are re-integrated after updating their list of variable genes, scaled, and reclustered (**Data File S1B**). Clusters containing cells with high mitochondrial, ribosomal, and non-coding RNAs are deemed to be low quality and are removed from our subsequent analyses. The clustering of Cd4 T cells is performed in the same manner (cluster markers in **Data File S1C**).

Uniform manifold approximation and projection (UMAP) plots, violin plots, dot plots, boxplots visualizations are done using functions from *Seurat* package or *ggplot2* package (version 3.5.1). Cell count frequency is calculated by normalizing cell count from each cluster to the total cell count of a specified cell type (e.g. Cd8 T cells or Cd4 T cells) in each sample. Pseudotime analysis is performed using Slingshot package (version 2.4.0). Heatmaps are generated using *ComplexHeatmap* package (2.12.1)

### Isolation of human CD8^+^ T cells

This study was conducted with approval from the institutional review boards of the University of California, Los Angeles (UCLA); written informed consent was obtained. All donors included in this study were female. For adult donors with T1D, blood was collected through UCLA endocrine clinics. For healthy adult donors, blood was collected through the UCLA Virology Core Laboratory. Collected blood was then performed peripheral blood mononuclear cell isolation, diluted with sterile PBS (Thermo Fisher Scientific). 35 mL of diluted blood was then overlayed with 15 mL of Ficoll-paque (GE Healthcare). The gradient was centrifuged at 400g with no brake for 30 min at room temperature. The PBMC interphase layer was collected, washed with FACS buffer, and centrifuged at 400g for 5 min, and cryopreserved in liquid nitrogen. For CD8^+^ T cell stimulation, PBMC CD8^+^ T cells were enriched with EasySep™ Human CD8^+^ T Cell Isolation Kit (STEMCELL Technologies, #17953).

### T cell stimulation

Mouse and human CD8^+^ T cells (1×10^6^/mL) were cultured in 10% FBS complete RPMI + human IL-2 (50 U/ml). Medium for mouse CD8^+^ T cells culture was supplemented with 1x 2-Mercaptoethanol (Gibco, 21985023). CD8^+^ T cells were plated in 48-well non-tissue culture plate for stimulation. Mouse CD8^+^ T cells were stimulated by Dynabeads™ Mouse T-Activator CD3/CD28, and human CD8^+^ T cells were stimulated by Dynabeads™ Human T-Activator CD3/CD28. After stimulation, mouse and human CD8^+^ T cells were analyzed by intracellular and surface protein staining. For TWS119 studies, 2×10^5^ of human CD8^+^ T cells were stimulated with Dynabeads® Human T-Activator CD3/CD28 under the condition with either DMSO or 5uM of TWS119 inhibitors to activate the Wnt/B-catenin pathway. After 3 days of stimulation, T_SCM_ and T_TE_ populations were identified by using flow cytometry with CD45RA and CCR7 markers.

### Genome editing in human resting CD8^+^ T cells by CRISPR

Freshly isolated human CD8^+^ T cells (2□×□10^6^) were washed twice with PBS and resuspended in 20□µl of buffer T (Thermo Fisher, Neon System). In parallel, synthetic sgRNAs (Synthego) were incubated with recombinant NLS-Cas9 (QB3 MacroLab) for 15□min at 37°C, at a ratio of 1:2.5 (40□pmol Cas9 protein per 100□pmol gRNA) to form the CRISPR–Cas9–gRNA RNP complex. CD8^+^ T cells were then added to the RNP complex and electroporated using the Neon NxT Electroporation System with the following settings: 2100 volts, 20 ms, and 1 pulse. Afterward, 1 mL of pre-warmed RPMI was added to the cells to allow recovery for 15 min at 37°C. Subsequently, complete RPMI medium supplemented with IL-7 (Peprotech; 200-07) and IL-15 (Peprotech; 200-15) (2 ng/ml each) was added. The sequence of sgRNA targeting the human KDM6A gene is as follows: UUGGAUAAUCUUCCAAUAAG and CAGCAUUAUCUGCAUACCAG. Scramble gRNA (Synthego) was used as a negative control. The sequences of the sgRNA used to target the STAT3 locus are GCAGCUUGACACACGGUACC and AAUGGAGCUGCGGCAGUUUC.

### RNA-seq library construction and data processing

Total RNA was isolated from sort-purified CD8^+^ progenitors (Ly108^+^ CD39^-^) using Quick RNA MiniPrep Kit (Zymo). RNA quality was verified using TapeStation analysis. RNA-seq libraries were generated by UCLA Technology Center for Genomics & Bioinformatics (TCGB) and sequenced on a Novaseq X Plus (paired-end, 2×50 bp). RNA-seq analysis was carried out by first checking the quality of the reads using FastQC. Then, they were mapped with HISAT2 (version 2.2.1) to the mouse genome (mm10). The counts for each gene were obtained by *featureCounts* package. Differential expression analyses were carried out using *DESeq2* (version 1.24.0) with default parameters. Genes with absolute fold change difference of > 1.5 with an FDR-adjusted P value□<□0.05 were considered significantly differentially expressed.

### CUT&Tag library preparation and data processing

CD8^+^ islet-specific progenitors (Ly108^+^ CD39^-^) was sorted by AIRA sorting and 300,000 progenitors were used as input for UTX CUT&Tag library preparation. After sorting, nuclei were isolated with cold nuclear extraction buffer (20□mM HEPES, pH 7.9, 10□mM KCl, 0.1% Triton X-100, 20% glycerol, 0.5□mM spermidine in 1× protease inhibitor buffer) and incubated with activated concanavalin A-coated magnetic beads (Polysciences, 86057-3) in PCR strip tubes at room temperature for 10□min. A 1:100 dilution of primary antibody (anti-UTX Cell Signaling Rabbit monoclonal antibody no. 33510 or IgG isotype control: Cell Signaling Technology, 3900S) in antibody buffer (20□mM HEPES pH 7.5; 150□mM NaCl, 0.5□mM spermidine, 1× protease inhibitor cocktail (Roche), 0.05% digitonin, 2□mM EDTA, 0.1% BSA) was added and nuclei were incubated with primary antibodies overnight at 4°C. The next day, the strip tubes were incubated on a magnetic tube holder and supernatants were discarded. Secondary antibody (guinea pig anti-rabbit IgG; Fisher Scientific, NBP172763) was added diluted at 1:100 in Dig-Wash (20□mM HEPES pH 7.5, 150□mM NaCl, 0.5□mM spermidine, 1× protease inhibitor cocktail, 0.05% digitonin) and nuclei were incubated for 1□h at room temperature. Nuclei were washed four times in Dig-Wash and then incubated with a 1:20 dilution of pAG–Tn5 adaptor complex (EpiCypher) in Dig-300 buffer (1× protease inhibitor cocktail, 20□mM HEPES pH 7.5, 300□mM NaCl, 0.5□mM spermidine) for 1□h at room temperature. To stop tagmentation, 25□μL Dig-300 buffer with 10□μL 1□M MgCl, 7.5□μL 0.5□M EDTA, 2.5□μL 10% SDS, and 5□μL 10□mg ml−1 proteinase K was added to each reaction and incubated at 55°C for 1□h. DNA was extracted by phenol:chloroform:isoamyl alcohol separation. DNA was barcoded and amplified using the following conditions: a PCR mix of 25□μL NEBNext 2× mix, 2□μL each of barcoded forward and reverse 10□μM primers, and 21□μL of extracted DNA was amplified at: 58°C for 5□min, 72°C for 5□min, 98°C for 45□s, 16× 98°C for 15□s followed by 63°C for 10□s, 72°C for 1□min. Amplified DNA libraries were purified by adding a 1.3× volume of KAPA pure SPRI beads (Roche) to each sample and incubating for 10□min at 23°C. Samples were placed on a magnet, and unbound liquid was removed. Beads were rinsed twice with 80% ethanol, and DNA was eluted with 25□μL TE buffer. All individually i7-barcoded libraries were mixed at equimolar proportions for sequencing on an Illumina Novaseq X Plus sequencer with 50bp pair-end sequencing.

The data analysis of UTX CUT&Tag, Stat3-ChIP (GEO: GSE217374), and TCF1-ChIP (GSE73240) was carried out from the raw fastq data file. The sequencing quality of libraries was assessed by *FastQC*. Ends of sequencing reads were trimmed by *Cutadapt*. *Bowtie2* (version 2.4.2) was used to align the sequencing reads to the mouse genome, mm10. PCR duplicates were removed using Picard *MarkDuplicates*. The uniquely mapped reads were used to call peaks with MACS2 using a p-value cutoff of 0.01. *ChIPseeker* was used for peak annotation. Figures of coverage tracks were exported from bigwig read alignment files using the UCSC Genome Browser.

### Co-Immunoprecipitation and Western blot

For the in vitro binding assay, 3.6×10^6^ of HEK-293T cells were plated on a 10-cm plate 24 h before transfection. Transfection was performed by mixing plasmids expressing Flag-tag UTX (Addgene plasmid #17438) and HA-Tag TCF1 (Addgene plasmid #40620) with the transfection reagent, TransIT-293 (Mirus), at a 1:2 (DNA: transfection reagent) ratio. For the STAT3 interaction, Flag-tag STAT3 (Addgene plasmid #8709) (51) or empty Flag construct was co-transfected with HA-UTX (Addgene plasmid #24168) (52). Twenty-four h post-transfection, cells were harvested and processed with Nuclear Complex Co-IP Kit (Active Motifs). 500 μg of total protein was used for co-immunoprecipitation, and Anti-FLAG® M2 Magnetic Beads were added to the samples for overnight incubation. After incubation, samples were washed 6 times with wash buffer and eluted in 2x LDS sample buffer, boiled at 95°C for 5 min. Pull-down samples were stored at −80 °C, and a western blot was performed for analysis.

For in vitro binding assay with Jurkat cells, 8×10^6^ Jurkat cells were used to isolate nuclear protein lysate with the Nuclear Complex Co-IP Kit. 500 µg of total protein was used for co-immunoprecipitation. 7 µg of UTX abs were added to pull down the protein complex, and rabbit IgG abs (7 µg) served as negative controls for overnight incubation at 4°C. After overnight incubation, 20 µL of Protein A (Cell Signaling Technology) for rabbit IgG was added to the protein lysate and incubated at 4°C for 1 h. After incubation, samples were washed six times with wash buffer and then eluted in 2× LDS sample buffer. They were subsequently boiled at 95°C for 5 min. Pull-down samples were stored at −80°C, and a western blot was performed for analysis.

Protein lysates were prepared in LDS sample buffer with 10 mM DTT. Samples were loaded and run onto precast gels, NuPAGE™ Bis-Tris Mini Protein Gels, 4–12%. Thereafter, gels were transferred to PVDF membrane with 10% methanol transferring buffer at 30 V for 60 min. Membranes were blocked in 5% skim milk in TBS-T buffer for 1 h at room temperature on a rocking platform. After blocking, membranes were incubated overnight with the following antibodies: HA-Tag (C29F4) Rabbit mAB (CST, # 3724) and Monoclonal ANTI-FLAG® M2 antibody (CST #8146 or #14793). Diluent and dilutions were determined following the manufacturer’s instructions. The following day, membranes were incubated with Goat anti-Rabbit IgG (H+L) Secondary Antibody, HRP (Abcam, # ab205718), or Anti-mouse IgG, HRP-linked Antibody (CST, # 7076), or infrared conjugated secondaries (IRdye 800CW Goat anti-Mouse IgG: LICOR #926-32210 or Goat anti-Rabbit 680LT: LICOR #926-68021) for 1 h at room temperature on a rocking platform.

### ATAC-seq library construction and data processing

CD8^+^ progenitors (Ly108^+^ CD39^-^) isolated from the pLN of *NOD-TCR 8.3 WT* (n=3) and *NOD-TCR 8.3 UTX^TCD^*(n=3) mice were purified by cell sorting. Each genotype has 3 biological replicates. 50,000 cells were used for ATAC-seq library preparation based on published Omni-ATAC protocol (53). In brief, the sorted cells were treated with lysis buffer for 3 min on ice, and the extracted nuclei were resuspended in the transposition mix containing 2.5 µL transposase (Illumina) and incubated at 37°C for 30 min with 1,000 rpm shaker. The products were purified with DNA Clean and Concentrator-5 Kit (Zymo Research) and then amplified with PCR for 11 cycles using barcoding primers provided by previous literature. PCR products were performed size selection using AMPure XP Bead (Beckman coulter) to purify DNA fragments from 150-1,000 bp. The concentration and quality control were measured by Qubit and Tapestation. The libraries were pooled based on molar concentrations and sequenced on Novaseq X Plus (paired-end, 2×50□bp).

The sequencing quality of the libraries was assessed by *FastQC* (0.11.9), followed by the trimming process using *Cutadapt* (4.6). *Bowtie2* (2.4.2) was used to align the sequencing reads to the mm10 mouse genome, and only uniquely mapped reads (MAPQ≥30) were retained for downstream. *SAMtools* (1.15.0) was used to convert SAM files into BAM files and sort BAM files. *Picard MarkDuplicates* (2.25) was used to remove duplicate reads in the BAM files. *MACS2* (2.2.9) was used for ATAC-seq peak calling (paired-end mode, FDR-adjusted q < 0.01). For differential binding analysis, we used *Diffbind* (3.20) package to determine differentially bound peaks found in all replicates for WT vs UTX^TCD^ with DESeq2 method. Peaks/regions identified as differentially accessible were annotated using the annotatepeaks.pl function from the HOMER analysis package. *FeatureCounts* Package (2.0.3) was used to count the number of reads that overlap each peak per sample. Chromatin regions were considered as significantly differentially accessible at a threshold of absolute log2 fold change difference in accessibility > 0.5 and FDR adjusted P < 0.05. Count-per-million-normalized bigwig files were generated for visualization using *Deeptools’* (3.5.6) BamCoverage function. Figures of coverage tracks were exported from bigwig read alignment files using the UCSC Genome Browser.

### Statistical analysis

Statistical analysis was performed using GraphPad Prism 9 or R for scRNA sequencing analysis. Unpaired two-tailed t tests were used to compare two groups while paired t tests were used for matched samples. For type 1 diabetic incidence curves, a log-rank (Mantel-Cox) test was used. Benjamini-Hochberg (also known as FDR method) adjusted p-values were reported for differentially expressed genes, for Pathway Analysis by Enrichr and gene set enrichment analysis by GSEA. Fold changes were calculated as (B-A)/A. A p-value of less than 0.05 was considered significant and data are presented as mean±SEM with the following symbols: *p<0.05, **p<0.01, *** p<0.001.

### Study approval

All experiments with mice were approved by the UCLA Animal Research Committee.

## Supporting information

Data file S1. Differential expression genes in each cluster of scRNA-seq dataset, related to STAR Methods-Single Cell RNA Analysis.

Data file S2. Differentially accessible peak_ATACseq.

Data file S3. Differentially expressed gene_RNAseq.

Data file S4. CD8+ T stem cell-like memory signature.

Data file S5. UTX_bound_peaks_CUT&TAG.

Supporting data values

Table S1. Antibodies used for flow cytometry and Western blot experiments.

## Data and materials availability

Bulk RNA sequencing (GEO: GSE282227), bulk ATAC sequencing (GEO: GSE282227), and single cell RNA sequencing (GEO: GSE283268) data were deposited in the National Center for Biotechnology Information Gene Expression Omnibus database. Values for all data points in the graphs are reported in the Supporting Data Values file. The paper did not include any original code. Software and packages are listed in the key resources table and method. All scripts of the bioinformatic analyses are available upon request from M.A.S.

## Author Contributions

H.C.C., M.G.L., W.L., and M.A.S. conceptualized the study. H.C.C., L.A.K., S.Z., Z.Z., and W.H. performed bioinformatics analysis. H.C.C., M.F.B., H.H.W., K.B.S., L.A.K., D.S., S.Z., E.C.M., S.S., C.L.H.P., X.Y., J.G.O., and N.M.W. performed the experiments. H.C.C., M.F.B., K.B.S., E.C.M., M.G.L., W.H., and M.A.S. contributed to data visualization. M.F.B., C.L.H.P., R.C., D.J.C, S.D.M, and C.M.R. consented T1D patient donors. M.A.S. wrote the original draft. M.F.B., M.G.L., W.H., and M.A.S. reviewed and edited the final manuscript.

## Acknowledgments

The authors thank the NIH Tetramer Core (Atlanta, GA) for providing the NRP-V7/H-2Kd PE tetramer. We also thank the Flow Cytometry Core at the University of California, Los Angeles for technical support and the Technology Center for Genomics & Bioinformatics (TCGB) at UCLA for the help of scRNA-, bulk RNA-seq, and ATAC-seq. We thank members of the UNC Animal Models Core and Director Dale O. Cowley for generating the NOD.UTX^DMD^ mice.

---

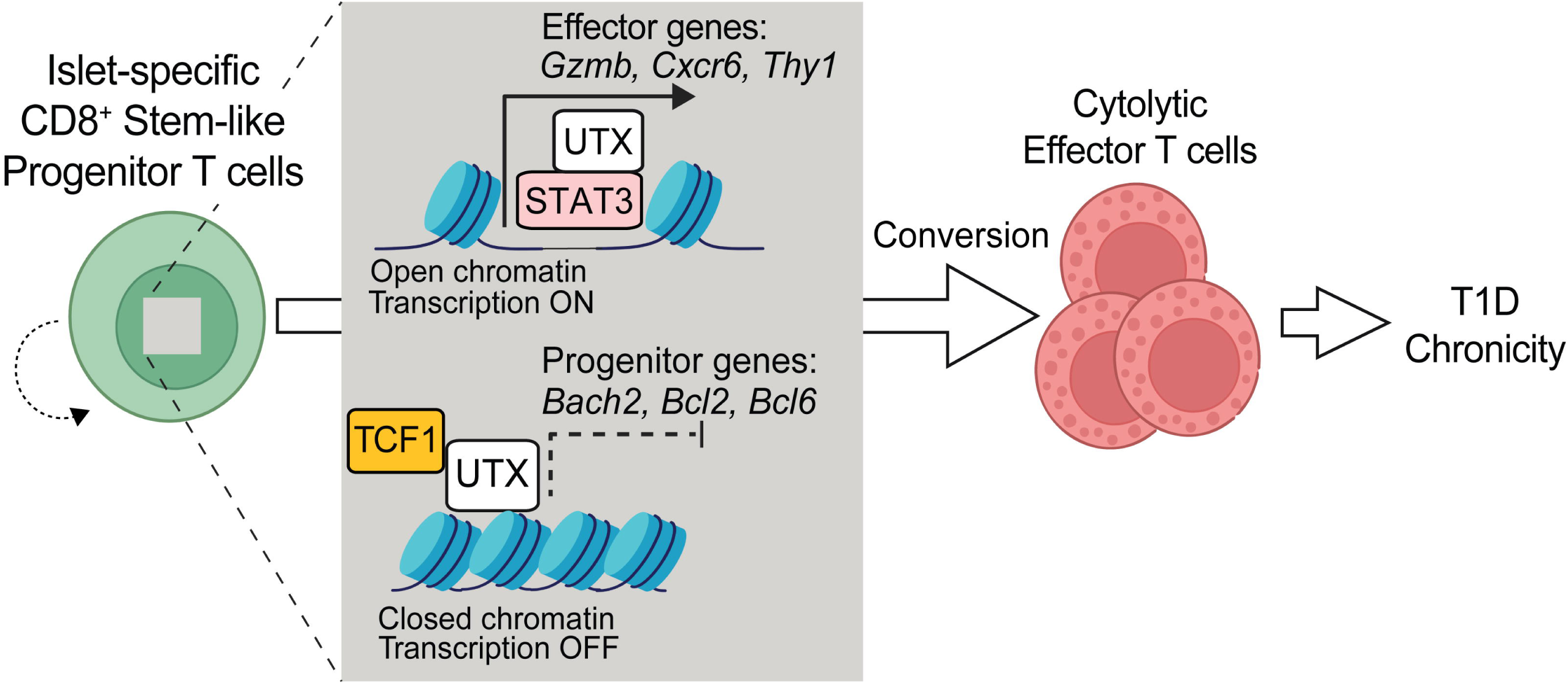

