## Supplemental Figures for "UTX coordinates TCF1 and STAT3 to control progenitor CD8^+^ T cell fate in autoimmune diabetes"

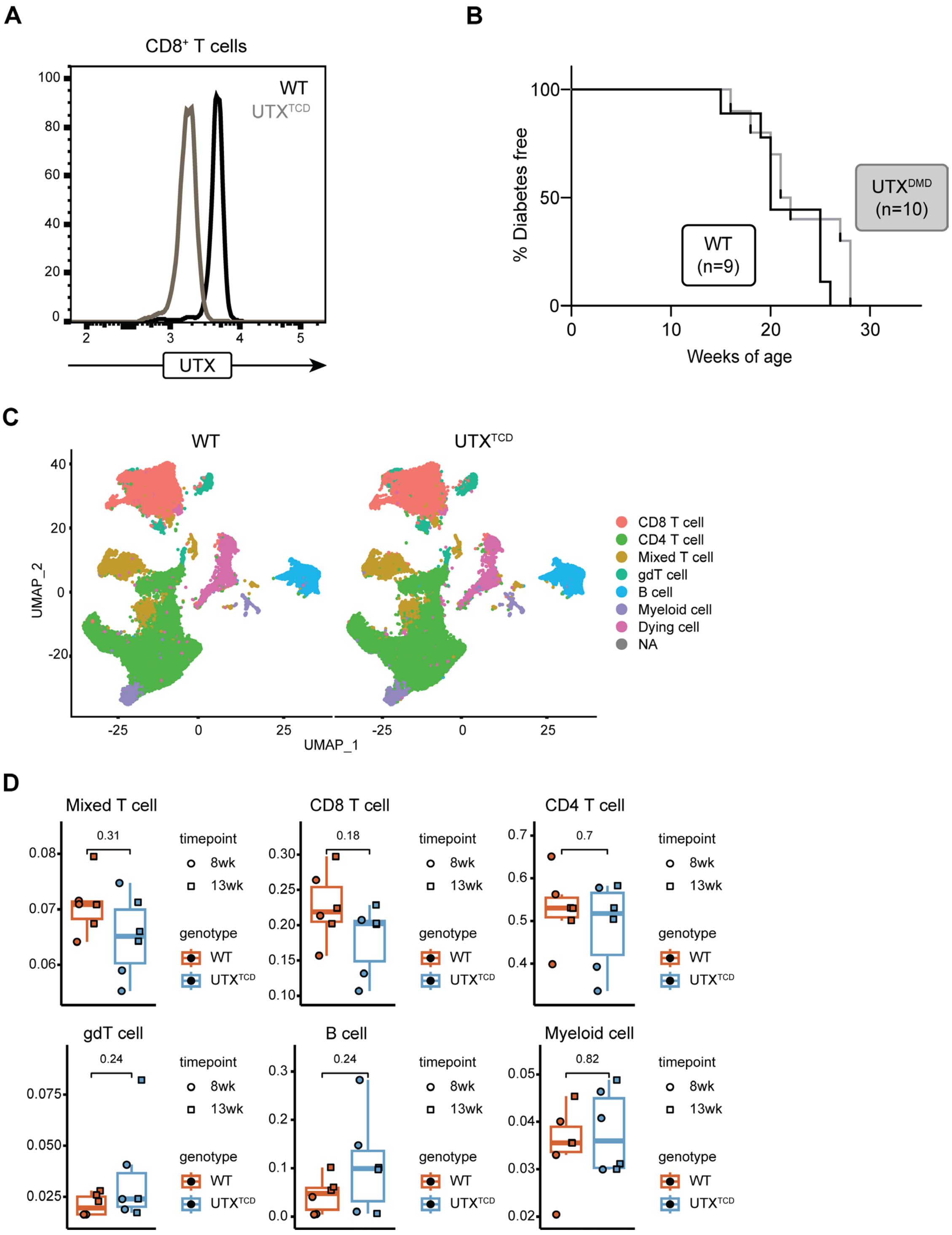


**Supplemental Figure 1. Single-cell RNA sequencing of CD45+ cell populations in *NOD-UTX$^{TCD}$* mice. (A)** Flow cytometry analysis on the UTX protein level of WT and UTX$^{TCD}$ CD8+ T cells. **(B)** Diabetes-free incidence curves of *NOD-UTX$^{DMD}$* vs. *NOD-WT* female littermates. ****p<0.0001; Log rank test. **(C)** Split UMAP of CD45+ cells by genotype. **(D)** Comparison of subset frequencies of CD45+ cell subsets. P values, two-sided, unpaired Mann-Whitney test.

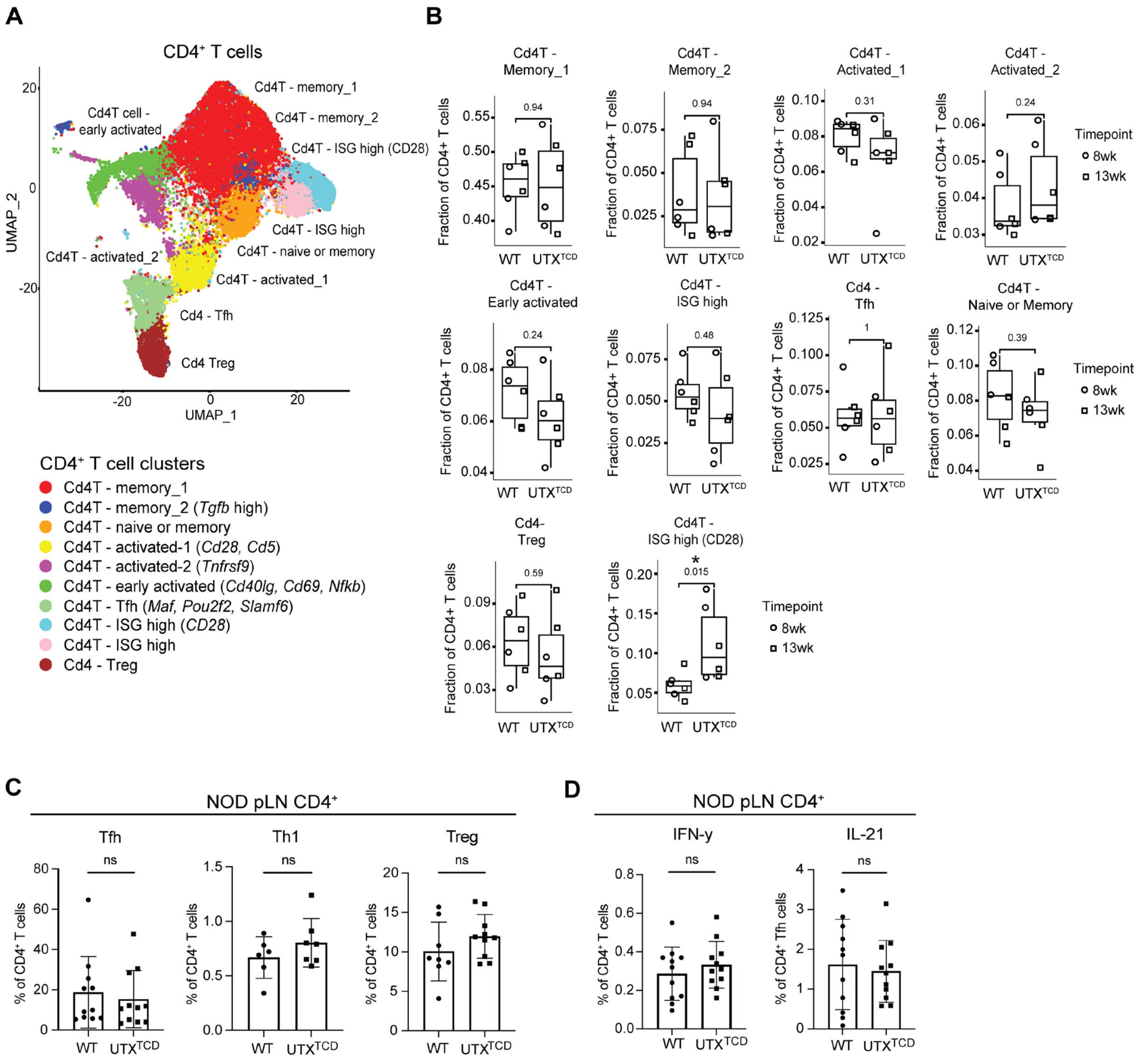


**Supplemental Figure 2. Minimal changes in CD4⁺ T cells from *NOD-UTX$^{TCD}$* mice. (A)** UMAP of CD4⁺ T cells. **(B)** Comparison of subset frequencies of CD4⁺ T cell subsets. P values, two-sided, unpaired Mann-Whitney test. **(C)** Average frequencies of Tfh (CXCR5⁺ PD1⁺), Th1 (T-bet⁺), and Treg (Foxp3⁺) cells among CD4⁺ T cells in pLN of *NOD-WT* and *NOD-UTX$^{TCD}$* female littermates. ns= not significant; unpaired Student's t-test. **(D)** Average frequencies of IFN-γ⁺ among CD4⁺ T cells and IL-21⁺ among CD4⁺ Tfh cells in pLN of *NOD-WT* and *NOD-UTX$^{TCD}$* female littermates. n.s.= not significant; unpaired Student's t-test.

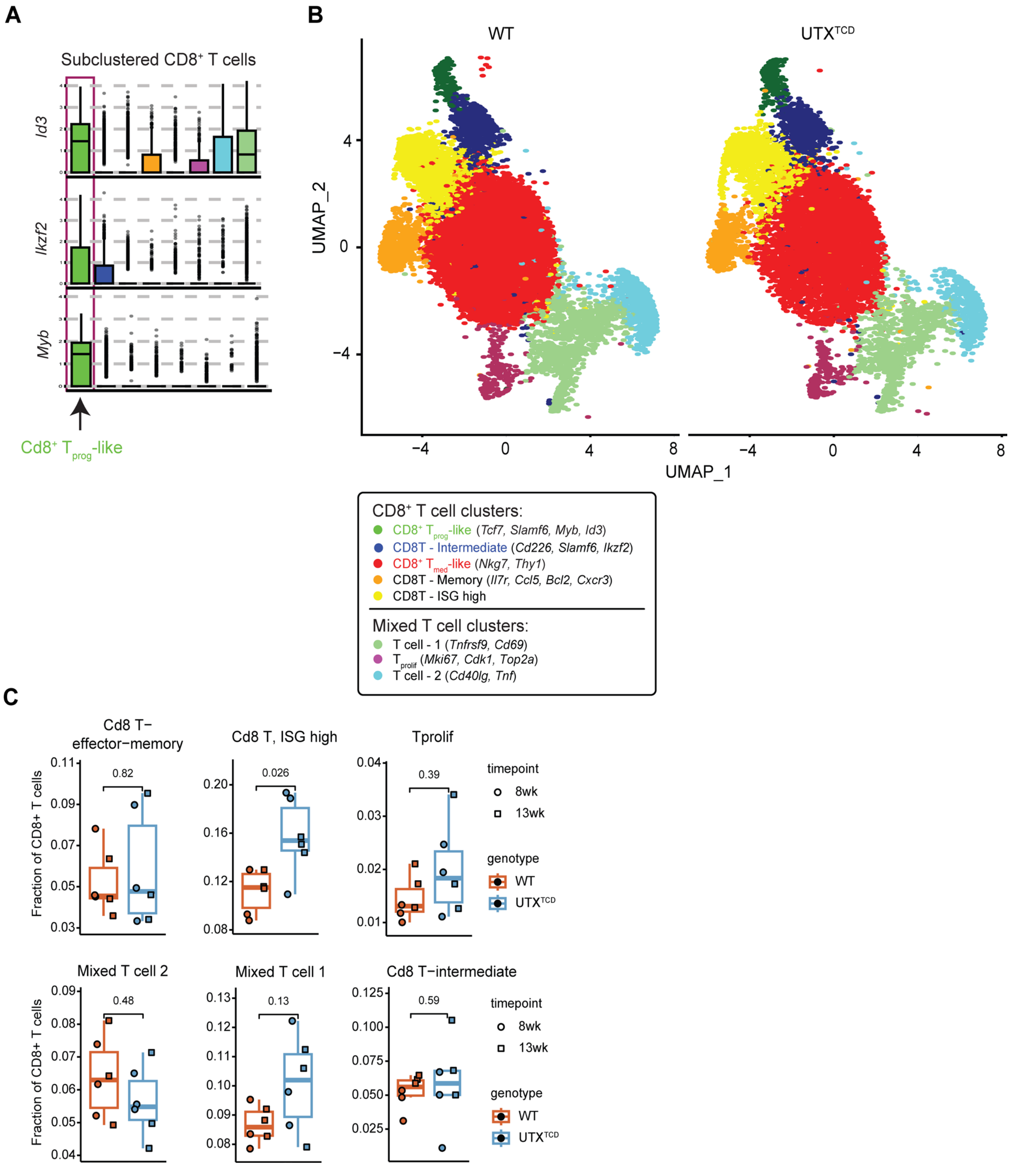


**Supplemental Figure 3. CD8$^{+}$ T cells from *NOD-UTX$^{TCD}$* mice. (A)** Expression of key genes (*Id3, Ikzf2,* and *Myb*) across CD8$^{+}$ T cell subclusters. **(B)** Split UMAP of CD8$^{+}$ T cells by genotype. **(C)** Comparison of subset frequencies of CD8$^{+}$ T cell subsets. Subset frequencies for “CD8 Tmed-like” and “CD8 Tprog-like” populations are shown in **Figure 1H**. P values, two-sided, unpaired Mann-Whitney test.

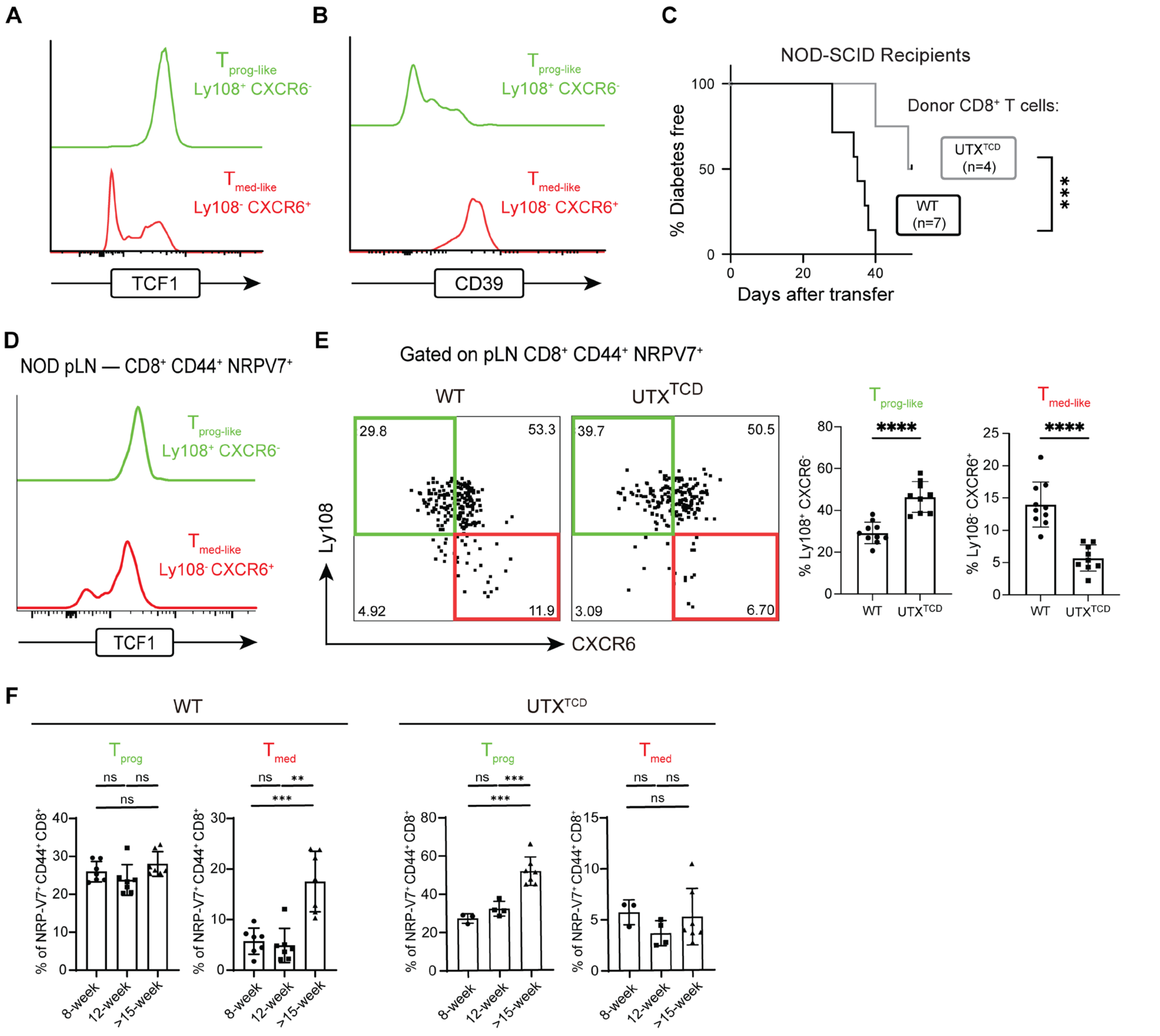


**Supplemental Figure 4. Altered CD8$^+$ T cell subset distribution in pLN of *NOD-UTX$^{TCD}$* mice.** **(A)** Representative histogram of TCF1 (stem-like marker) and **(B)** CD39 (effector marker) expression in $T_{prog}$-like vs. $T_{med}$-like cell populations. **(C)** Diabetes incidence curves of *NOD-SCID* mice recipients transferred with WT CD4$^+$ T cells and either *NOD-WT* CD8$^+$ T cells or *NOD-UTX$^{TCD}$* CD8$^+$ T cells. ***p<0.001; Log rank test. **(D)** Representative histogram of TCF1 (stem-like marker) in $T_{prog}$-like (Ly108$^+$ CD39$^-$) vs. $T_{med}$-like (Ly108$^-$ CD39$^+$) in pancreatic lymph nodes. **(E)** Representative flow cytometric plot (left) and average frequencies (right) of $T_{prog}$-like (Ly108$^+$ CXCR6$^-$) and $T_{med}$-like (Ly108$^-$ CXCR6$^+$) cells within antigen-experienced, IGRP-specific pLN CD8$^+$ T cells (CD8$^+$ CD44$^+$ NRPV7$^+$) of *NOD-WT* and *NOD-UTX$^{TCD}$* female littermates (12-16 weeks of age). ***p<0.001; Student's t test. **(F)** The frequency of IGRP$^+$ $T_{prog}$ (Ly108$^+$ CD39$^-$) and $T_{med}$ (Ly108$^-$ CD39$^+$) population in 8-week-old, 12-week-old, and over 15-week-old *NOD-WT* and *NOD-UTX$^{TCD}$* female mice. ***p<0.001; **p<0.01; unpaired two-way Mann Whitney test.

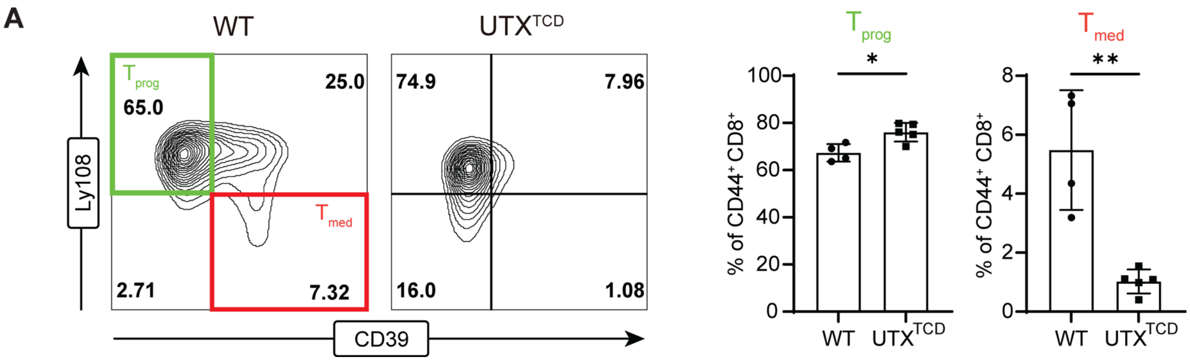


**Supplemental Figure 5. The phenotype of CD8$^{+}$ T cells in the pancreas of *NOD-TCR 8.3 UTX$^{TCD}$* mice. (A)** Representative flow plots (left) and frequencies (right) of $T_{prog}$ and $T_{med}$ in the pancreas of *NOD-TCR 8.3 WT* and *NOD-TCR 8.3 UTX$^{TCD}$* mice. **p<0.01; *p<0.05; Student's t test.

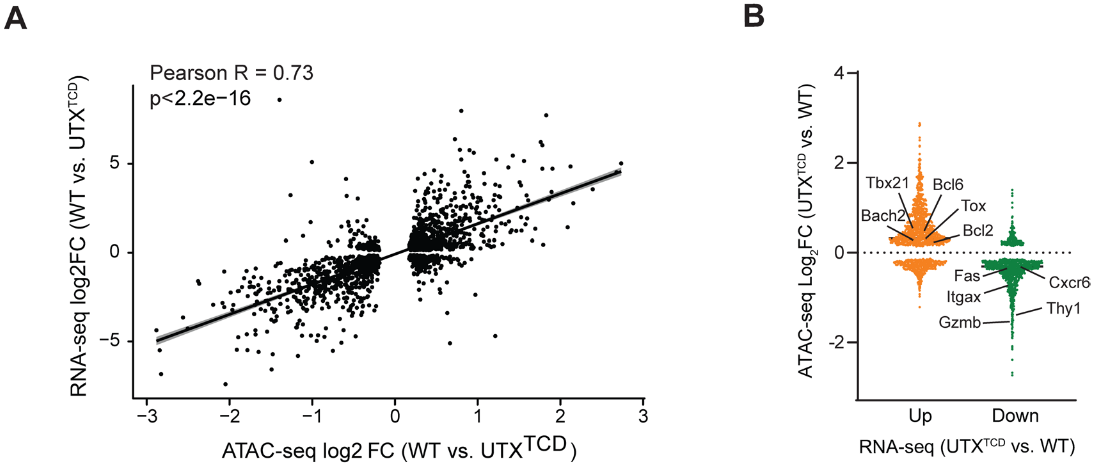


**Supplemental Figure 6. Analysis of ATAC-seq data and RNA-seq reveals differentially regulated genes. (A)** Correlation between the differential chromatin accessibility and differential gene expression values (in log2FC) in the CD8$^+$ $T_{prog}$ cells of *NOD-TCR 8.3 UTX$^{TCD}$* compared to *NOD-TCR 8.3 WT* (difference in RNA expression, y-axis; difference in ATAC accessibility, x-axis). R, Pearson correlation coefficient. **(B)** Difference in chromatin accessibility of listed genes (in log2FC) between *NOD-TCR 8.3 UTX$^{TCD}$* and *NOD-TCR 8.3 WT* CD8$^+$ $T_{prog}$ cells as measured by ATAC-seq (decreased accessibility in *NOD-TCR 8.3 UTX$^{TCD}$*, log2 FC < -0.5; increased in *NOD-TCR 8.3 UTX$^{TCD}$*, log2 FC > 0.5).

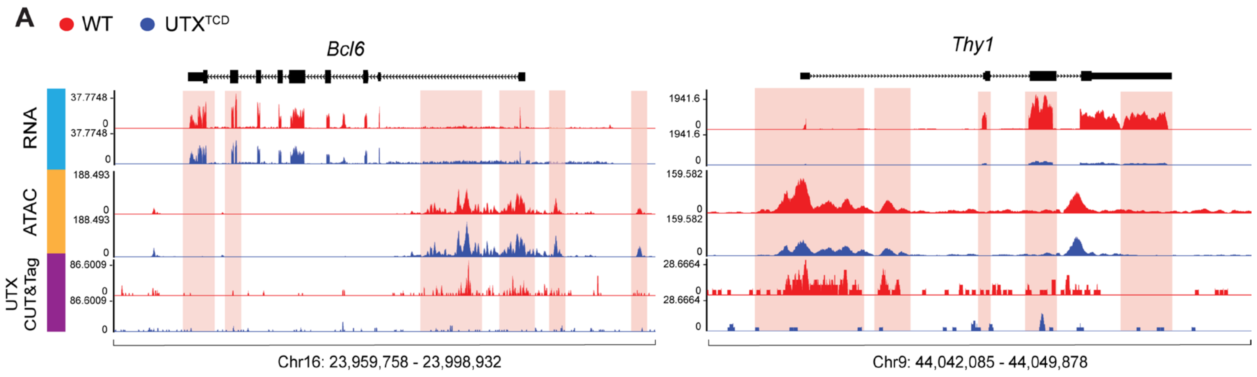


**Supplemental Figure 7. UTX deficiency alters chromatin accessibility, gene expression, and associated pathways. (A)** Representative gene tracks from UCSC Integrated Genome Browser of UTX CUT&Tag, ATAC-seq, and RNA-seq at Bcl6, and Thy1 gene loci.

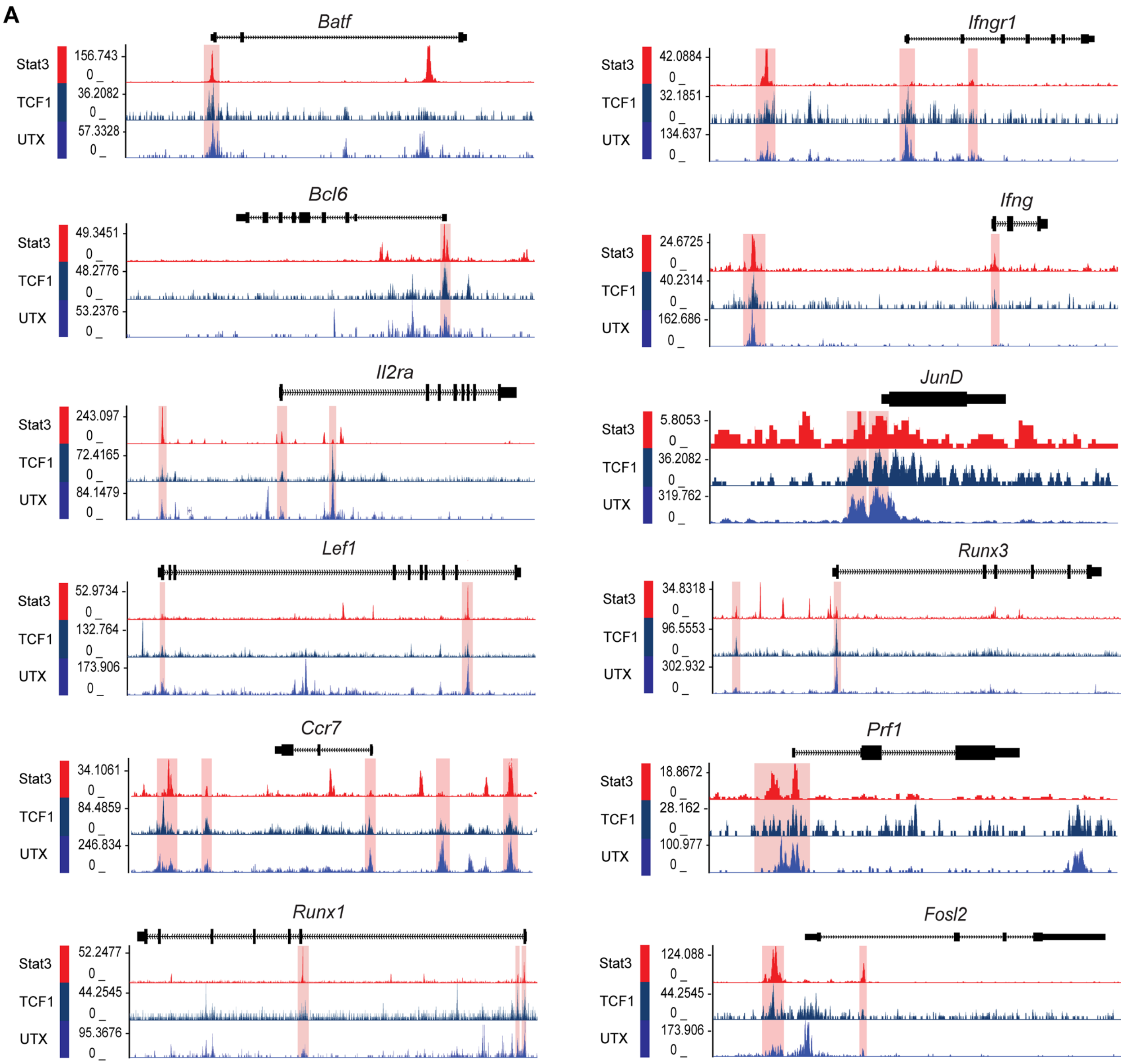


**Supplemental Figure 8. Co-occupancy of STAT3, TCF1, and UTX at progenitor and effector genes. (A)** Representative gene tracks from the UCSC genome browser to demonstrate the aligned region bound by UTX, TCF1, and Stat3 in progenitor genes (*Batf, Bcl6, Il2ra, Lef1, Ccr7,* and Runx1; left) and mediator genes (*Ifngr1, Ifng, JunD, Runx3, Prf1,* and *Fosl2*; right).

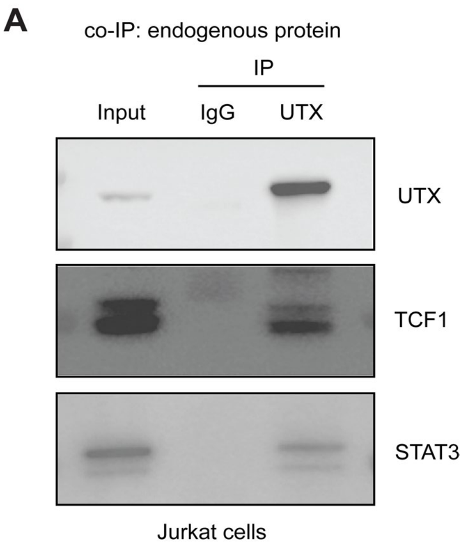


**Supplemental Figure 9. UTX interacts with TCF1 and STAT3 in Jurkat T cells. (A)** UTX co-immunoprecipitation (co-IP) of endogenous proteins from Jurkat cells. Immunoprecipitants were analyzed by immunoblotting for UTX, TCF1, and STAT3. Input and IgG pulldown controls are shown.
