## Supplementary material for "UTX coordinates TCF1 and STAT3 to control progenitor CD8^+^ T cell fate in autoimmune diabetes": Table S1. Antibodies used for flow cytometry and Western blot experiments.

| | | |
|---|---|---|
| Brilliant Violet 711™ anti-mouse CD8a Antibody | Biolegend | Cat# 100748;<br>RRID: AB_2562100 |
| APC anti-mouse Ly108 Antibody | Biolegend | Cat# 134610;<br>RRID: AB_2728155 |
| FITC anti-mouse CD186 (CXCR6) Antibody | Biolegend | Cat# 151108;<br>RRID: AB_2572145 |
| APC/Cyanine7 anti-mouse CD3 Antibody | Biolegend | Cat# 100222;<br>RRID: AB_2242784 |
| Alexa Fluor® 700 anti-mouse CD45 Antibody | Biolegend | Cat# 103128;<br>RRID: AB_493715 |
| PE/Dazzle™ 594 anti-mouse/human CD44 Antibody | Biolegend | Cat# 103056;<br>RRID: AB_2564044 |
| BD Horizon™ BUV395 Rat Anti-Mouse CD39 | BD<br>Biosciences | Cat# 567264;<br>RRID: AB_2916524 |
| CD279 (PD-1) Monoclonal Antibody (J43), PE,<br>eBioscience™ | Thermo Fisher<br>Scientific | Cat# 12-9985-82;<br>RRID: AB_466295 |
| PE/Cyanine7 anti-mouse CD279 (PD-1) Antibody | Biolegend | Cat# 109110;<br>RRID: AB_572017 |
| PE/Cyanine7 anti-mouse CD185 (CXCR5) Antibody | Biolegend | Cat# 145516;<br>RRID: AB_2562210 |
| FITC anti-mouse CD185 (CXCR5) Antibody | Biolegend | Cat# 145520;<br>RRID: AB_2562866 |
| Brilliant Violet 605™ anti-mouse CD4 Antibody | Biolegend | Cat# 100548;<br>RRID: AB_2563054 |

| | | |
|---|---|---|
| IL-21 Monoclonal Antibody (mhalx21), PE, eBioscience™ | Thermo Fisher Scientific | Cat# 12-7213-82; RRID: AB_1834465 |
| APC anti-mouse IFN-γ Antibody | Biolegend | Cat# 505810; RRID: AB_315404 |
| PE anti-T-bet Antibody | Biolegend | Cat# 644810; RRID: AB_2200542 |
| FOXP3 Monoclonal Antibody (FJK-16s), FITC, eBioscience™ | Thermo Fisher Scientific | Cat# 11-5773-82; RRID: AB_465243 |
| Brilliant Violet 711™ anti-human CD8 Antibody | Biolegend | Cat# 344734; RRID: AB_2565243 |
| PE anti-human CD4 Antibody | Biolegend | Cat# 300550; RRID: AB_2564152 |
| PE/Cyanine7 anti-human CD95 (Fas) Antibody | Biolegend | Cat# 305622; RRID: AB_2100369 |
| FITC anti-human CD45RA Antibody | Biolegend | Cat# 304148; RRID: AB_2564157 |
| Brilliant Violet 605™ anti-human CD45RO Antibody | Biolegend | Cat# 304238; RRID: AB_2562153 |
| APC/Cyanine7 anti-human CD197 (CCR7) Antibody | Biolegend | Cat# 353212; RRID: AB_10916390 |
| TCF1/TCF7 (C63D9) Rabbit mAb (PE-Cy7® Conjugate) | Cell Signaling Technology | Cat# 90511S; RRID: AB_3086656 |
| TCF1/TCF7 (C63D9) Rabbit mAb (APC Conjugate) | Cell Signaling Technology | Cat# 37636S; RRID: AB_2922379 |

| | | |
|---|---|---|
| KDM6A antibody [N2C1], Internal | GeneTex | Cat# GTX121246; RRID:AB_10722382 |
| InVivoMAb anti-mouse PD-1 (CD279) | BioXCell | Cat# BE0146; RRID:AB_10949053 |
| InVivoMAb mouse IgG2b isotype control | BioXCell | Cat# BE0086; RRID: AB_1107791 |
| Goat Anti-Rabbit IgG H&L (FITC) | abcam | Cat# ab6717; RRID:AB_955238 |
| Tri-Methyl-Histone H3 (Lys27) (C36B11) Rabbit mAb | Cell Signaling Technology | Cat# 9733S; RRID:AB_2616029 |
| UTX (D3Q1I) Rabbit mAb | Cell Signaling Technology | Cat# 9733S; RRID:AB_2721244 |
| DYKDDDDK Tag (D6W5B) Rabbit mAb | Cell Signaling Technology | Cat# 9733S; RRID:AB_2572291 |
| HA-Tag (C29F4) Rabbit mAb | Cell Signaling Technology | Cat# 9733S; RRID:AB_1549585 |
